# An Alternative DNA Endonuclease Activity is Associated with the LINE-1 ORF2-encoded Protein

**DOI:** 10.64898/2026.01.20.700639

**Authors:** Mitsuhiro Nakamura, Huira C. Kopera, Mark Dowling, Orsolya Barabas, John V. Moran

## Abstract

Long INterspersed Element-1 (L1) retrotransposons use activities contained within the L1 open reading frame 2-encoded protein (ORF2p) to mobilize throughout the genome via target-site primed reverse transcription (TPRT). The ORF2p endonuclease domain (EN) cleaves genomic DNA to liberate a 3’-hydroxyl (3’-OH) group that is used by the ORF2p reverse transcriptase domain (RT) to synthesize a cDNA copy of its bound L1 RNA template. L1 also can move by EN-independent retrotransposition (ENi), where a 3’-OH group at genomic DNA lesions, dysfunctional telomeres, or stalled replication forks is proposed to prime L1 reverse transcription in the absence of L1 EN cleavage. We previously reported that ribonucleoprotein (RNP) preparations from cells transfected with a human wild-type (WT) L1 or L1 EN-mutant, but not an L1 RT-mutant, can initiate reverse transcription from a DNA oligonucleotide primer/L1 RNA template complex. The WT and EN-deficient L1 RNP preparations also are associated with a nuclease activity that can process a 3’ end modification that precludes DNA synthesis from an oligonucleotide prior to priming the L1 RT reaction. Here, we purified recombinant full-length WT, L1 EN-, and L1 RT-mutant human L1 ORF2p from insect cells. We report that the WT and L1 EN-mutant, but not the L1 RT-mutant, contain an alternative endonuclease activity (alt-EN). Alt-EN activity also is detected in a bacterially expressed L1 ORF2p protein that lacks the L1 EN and ORF2p cysteine-rich domains and a thermostable group II intron-encoded protein. Processing of diverse modified primers demonstrates endonucleolytic cleavage that is eliminated by mutations in the RT active site. We propose that alt-EN is an evolutionarily conserved activity within the RT fold that promoted ENi retrotransposition of primordial retrotransposons prior to the acquisition of an EN domain.

## Introduction

Long Interspersed Element-1 (LINE-1 or L1) is a non-long terminal repeat (non-LTR) retrotransposon that can mobilize to new genomic locations via a “copy and paste” mechanism termed retrotransposition (1). Although L1-derived sequences are present at greater than 500,000 copies in the human genome (2), an average individual genome only contains ∼100-150 retrotransposition competent L1s (RC-L1s) (3–7). RC-L1s are approximately six kilobases in length and encode a sense strand bicistronic mRNA containing two open reading frames (*L1 ORF1* and *L1 ORF2*) (8,9). *L1 ORF1* encodes a 40 kDa protein, ORF1p, with RNA binding and nucleic acid chaperone activities (10–18). *L1 ORF2* encodes a 150 kDa protein, ORF2p, with endonuclease (EN) and reverse transcriptase (RT) activities (17,19–24). ORF2p also has a conserved carboxyl-terminal cysteine rich RNA binding domain (C) (17,25). Mutational analyses revealed that conserved amino acids within ORF1p and ORF2p are required for retrotransposition (11,17,21).

Canonical L1 retrotransposition occurs via target-site primed reverse transcription (TPRT) (17,18,21,26–30). A full-length human RC-L1 mRNA is transcribed from an internal RNA polymerase II promoter present within its 5’ untranslated region (UTR) (18,31–35). The full-length bicistronic L1 mRNA then is exported to the cytoplasm where it undergoes translation (19,20,36–38). Abundant experimental evidence demonstrates that ORF1p and ORF2p bind to their encoding RNA in *cis* (which we termed *cis-*preference (39)), forming a ribonucleoprotein (RNP) particle that minimally consists of one molecule of full-length L1 mRNA, multiple molecules of ORF1p, and one molecule of ORF2p (11,18,19,39–41). Components of the L1 RNP enter the nucleus to initiate TPRT (42,43). The ORF2p EN domain generates a single-strand endonucleolytic nick in double-stranded genomic DNA at a degenerate consensus sequence (*e.g.*, 5’-TTTTT/AA-3’ and variants of that sequence, where “/” indicates the cleavage position), liberating a 3’-hydroxyl (OH) group that is used as a primer by the ORF2p RT domain to generate a cDNA copy of its bound L1 mRNA (17,21,27,29,44–48). Although subsequent steps of L1 integration require elucidation (*i.e.*, second strand cleavage and second strand cDNA synthesis), a union of previous and emerging experimental evidence suggests that ORF2p and/or host proteins can facilitate the completion of TPRT, leading to the integration of an L1 copy at a new genomic location (18,26,28,30,49,50). The new L1 insertion has the following structural hallmarks: (**1**) it ends in a 3’ poly(A) tract; (**2**) it is located at an L1 EN cleavage site; and (**3**) it generally is flanked by short target site duplications that range in size from ∼7-20 bp (reviewed in (18,51)).

Our previous studies discovered that L1s can exploit lesions in genomic DNA, thereby bypassing the requirement for L1 EN DNA cleavage. This mechanism, which we termed L1-EN independent retrotransposition (ENi) (52), is supported by several lines of experimental evidence. First, EN-deficient L1s can retrotranspose in Chinese hamster ovary (CHO) cells defective for p53 and components of the non-homologous end-joining (NHEJ) pathway of DNA repair and human HCT116 colorectal cancer cells that are both p53- and DNA ligase IV-deficient (52,53). Second, EN-mutant L1s can integrate at dysfunctional telomeres in certain p53-/NHEJ-deficient CHO cells (54,55). Third, recent data indicate that an L1 EN-mutant can retrotranspose efficiently in a FANCD2-deficient human cell line, presumably by initiating L1 cDNA synthesis from a 3’-OH group present at an Okazaki fragment (45). These data strongly suggest that the L1 RT can use a 3’-OH group at endogenous DNA lesions to prime the reverse transcription of L1 mRNA. Thus, we proposed that ENi represents an ancestral type of RNA-mediated DNA repair that was used by LINE elements to mobilize before the acquisition on an endonuclease domain (45,52).

We developed an *in vitro* assay, termed the L1 element amplification protocol (LEAP), which allows the detection of L1 RT activity in RNP preparations derived from cells transfected with either wild type (WT) or mutant human L1 constructs (11,41,56). Briefly, RNPs derived from cells transfected with engineered human L1s are incubated with an oligonucleotide primer ending in a 3’ poly(T) tract. This oligonucleotide can anneal with the poly(A) tails of mRNAs present within the preparation; however, only RNAs bound by the L1 RT in *cis* template cDNA synthesis within the oligonucleotide primer/L1 mRNA complex. PCR amplification of the resultant L1 cDNAs then generates a robust readout of L1 RT activity (**Fig. S1**). The use of the LEAP assay revealed several important aspects of L1 biology and ORF2p function: (**1**) WT and EN-deficient L1 RNPs, but not RT-deficient L1 RNPs, exhibit LEAP activity (41); (**2**) terminal complementarity between the oligonucleotide primer and the L1 mRNA template is not required by the L1 RT to initiate the reverse transcription of L1 mRNA (41,57); and (**3**) a functional version of ORF1p is not required to initiate the reverse transcription of L1 mRNA, but helps position the initiation of L1 cDNA synthesis within the L1 3’ poly(A) tail (11,41,56).

We observed that L1 RNP preparations also contain a nuclease activity that could remove 3’ modifications that preclude DNA synthesis from an oligonucleotide primer, allowing it to be used to prime L1 cDNA synthesis (54). However, because L1 RNP preparations contain host proteins and host RNAs, we could not determine whether ORF2p or a cellular protein was responsible for this nuclease activity. Here, we used a partially purified full-length version of ORF2p immunoprecipitated from human embryonic kidney (HEK293T) cells (IP-ORF2p) (58), and a full-length recombinant version of (rORF2p) purified from insect cells, in a modified LEAP assay and an RT-independent ligation assay to demonstrate that ORF2p itself possesses an alternative endonuclease activity (herein called alt-EN). Importantly, we show that alt-EN cleaves internal DNA sugar-phosphate bonds, and a missense mutation in the ORF2p RT active site (D702A) severely dimishes this activity. Moreover, we demonstrate that an ORF2p “core” protein purified from bacteria that lacks both the L1 EN domain and carboxyl-terminal cysteine-rich (C) domain (26), as well as a commercially available thermostable group II intron-encoded RT protein, contain alt-EN activity. Thus, we propose that alt-EN is an evolutionarily conserved enzymatic activity associated with the RT domain of non-LTR retrotransposons that can remove a 3’ modification that blocks DNA synthesis from lesions in genomic DNA to facilitate ENi retrotransposition.

## Materials and Methods

### Cell culture

HeLa-JVM and HEK293T cells were grown in high-glucose DMEM (Invitrogen) medium supplemented with 10% FBS, 100 U/ml penicillin-streptomycin, and 0.29 mg/ml L-glutamine (DMEM-complete medium) at 37°C in a 7% CO_2_ humidified tissue culture incubator. *Spodoptera frugiperda* 9 (*Sf*9) cells were grown in Sf-900 II SFM (Gibco), Sf-900 III SFM (Gibco), or Insect-Xpress (Lonza) medium at 27°C with shaking at 120 rpm. High Five cells were grown in EXPRESS FIVE SFM (Gibco) or Insect-Xpress medium at 27°C with shaking at 120 rpm.

### Oligonucleotides

DNA and RNA oligonucleotides used in this study in the Moran laboratory were purchased from Integrated DNA Technologies (IDT). DNA oligonucleotides used in the Barabas laboratory were purchased from Microsynth (MN462 and MN085) or IDT (4228 and MN503). The names and sequences of the oligonucleotides are listed in **Table S1**. The oligonucleotide primers used to prime cDNA synthesis during the reverse transcription steps in LEAP and modified LEAP assays (41,56) and the substrate oligonucleotides used in the RT-independent ligation-based assay to monitor alt-EN activity were purified by IDT using high pressure liquid chromatography (HPLC). DNA oligonucleotides purchased from Microsynth also were purified using HPLC.

### Plasmids

All the L1 expression plasmids used in this study contain the L1.3 DNA sequence (GenBank accession no. L19088) (6). For mutant plasmids, the amino acid residues were counted from the first methionine (M1) of the L1.3 ORF2p sequence. Plasmid DNAs used in human cell transfection experiments were purified using a Qiagen Plasmid Midi Kit. Other plasmid DNAs used in the study were purified using a Wizard Plus SV Minipreps DNA Purification System (Promega).

#### The following plasmids were used in this study

pCEP4: is a commercially available mammalian episomal expression vector (Invitrogen). It was used to express WT and mutant L1.3 sequences in cultured human cells.

pTMO2F3: was described previously (59). The plasmid contains the L1.3 5’UTR, a copy of the *L1.3 ORF2* sequence containing three tandem copies of an in-frame FLAG epitope-tag (3xFLAG) sequence at the 3’ end of *ORF2*, and a partial sequence of the L1.3 3’UTR; it is cloned into the pCEP4 expression plasmid (Invitrogen). A cytomegalovirus immediate early (CMV) promoter and simian vacuolating virus 40 (SV40) polyadenylation signal in the pCEP4 plasmid augment the expression of *L1.3 ORF2-3xFLAG*.

pAD001: was described previously (59). It is similar to pTMO2F3 but lacks a 3xFLAG epitope-tag sequence.

pTMO2F3_D145A: was described previously (58). It is identical to pTOM2F3 but contains a D145A missense mutation in the L1 ORF2p EN domain (39).

pTMO2F3_D205A: was described previously (58). It is identical to pTMO2F3 but contains a D205A missense mutation in the L1 ORF2p EN domain (39).

pTMO2F3_D702A: was described previously (58). It is identical to pTMO2F3 but contains a D702A missense mutation in the L1 ORF2p RT domain (39).

pMN001: contains a copy of *L1.3 ORF2* containing three tandem copies of an in-frame human influenza hemagglutinin (HA) epitope-tag (3xHA) and a 3xFLAG epitope tag sequence at the 3’ end of *ORF2*. A polyhedrin promoter and SV40 polyadenylation signal in the pFastBac1 plasmid (Invitrogen) augment the expression of *L1.3 ORF2-3xHA/3xFLAG*.

pMN001_D205A: is identical to pMN001 but contains a D205A missense mutation in the L1 ORF2p EN domain (39).

pMN001_D702A: is identical to pMN001 but contains a D702A missense mutation in the L1 ORF2p RT domain (39).

pJJ101/L1.3: was described previously (54). It contains a full-length copy of L1.3 (L1.3 5’UTR, *L1.3 ORF1*, *L1.3 ORF2*, and L1 3’UTR). The L1 3’UTR contains a blasticidin (*mblastI*) retrotransposition indicator cassette. A CMV immediate early promoter and SV40 polyadenylation signal in the pCEP4 plasmid augment L1.3 expression.

pJJ105/L1.3: was described previously (54). It is identical to pJJ101/L1.3 but contains a D702A missense mutation in the L1.3 RT ORF2p domain (39).

pJJD145A/L1.3: was described previously (54). It is identical to pJJ101/L1.3 but contains a D145A missense mutation in the L1.3 ORF2p EN domain (39).

pJJD205A/L1.3: was described previously (54). It is identical to pJJ101/L1.3 but contains a D205A missense mutation in the L1.3 ORF2p EN domain (39).

pAJ101/L1.3: was described previously (54). It is similar to pJJ101/L1.3 but lacks the *L1.3 ORF1* sequence.

pT4Zf1: is a plasmid that was used to generate a standard curve for the quantitative PCR (qPCR) reaction described in the RT-independent ligation-based assay section (see below). Briefly, the MN439 and MN544 oligonucleotides (**Table S1**) were ligated; the product then was used as a template in a PCR reaction containing the MN547 and MN549 oligonucleotides (**Table S1**). The resultant PCR product was cloned into the pCR_Blunt vector (Invitrogen). Known concentrations of the resultant DNAs were used as a control standard in the RT-independent ligation-based assay section (also see below).

pJCC5/L1.3: was described previously (17). It contains a full-length copy of L1.3 (L1.3 5’UTR, *L1.3 ORF1*, *L1.3 ORF2*, and L1 3’UTR) and is cloned into the pBLS KS-vector (Stratagene).

p1253_ORF2p-StrepII: is identical to pMN001 but the 3xHA epitope-tag and 3xFLAG epitope-tag sequence is replaced by a GGS linker sequence followed by a Strep-Tag II sequence at the 3’ end of *ORF2*.

p1393_ORF2p-D702A-StrepII: is identical to p1253_ORF2p-StrepII but contains a D702A missense mutation in the L1 ORF2p RT domain.

### Immunoprecipitation of L1 ORF2p-3xFLAG tag

The immunoprecipitation of L1.3 ORF2p-3xFLAG from transfected human HEK293T cells was modified slightly from a previously described protocol (58). HEK293T cells were plated in a T175 flask (BD Biosciences) containing DMEM-complete medium at approximately 5 x 10^6^ cells per flask. The following day (day 0), the cells were transfected with 30 µg of the indicated plasmids (*i.e.*, pCEP4, pAD001, pTMO2F3, pTMO2F3_D145A, pTMO2F3_D205A, and pTMO2F3_D702A) using 90 µL of the FuGENE6 transfection reagent (Promega) and 3 mL of Opti-MEM (Gibco) following the protocol provided by the manufacturer. The following day (day 1), the transfection medium was replaced with fresh DMEM-complete medium. On days 2-5, the medium was replaced daily with DMEM-complete medium containing 200 µg/mL hygromycin B (Gibco). On day 5, the cells were collected, washed with cold 1x phosphate buffered saline solution (PBS, Gibco), and flash-frozen in a dry ice/ethanol bath.

The subsequent steps in the protocol were performed on ice or at 4°C unless noted otherwise. One mL of Lysis300 buffer (20 mM Tris-HCl pH 7.5, 2.5 mM MgCl_2_, 1 mM dithiothreitol [DTT], 300 mM KCl, and 0.1% IGEPAL CA-630) containing 1x cOmplete EDTA-free Protease Inhibitor Cocktail (Roche) and 0.2 mM phenylmethylsulfonyl fluoride (PMSF) was added to the transfected cells. The cells then were incubated for one hour with gentle rotation. To obtain whole-cell lysates (WCLs), the insoluble cell debris from the resultant solution was removed by centrifugation at 15,000 x *g* for five minutes. The protein concentration of the resultant WCLs was determined using the Bradford reagent assay (BioRad).

Prior to immunoprecipitation, antibody-attached beads were prepared by incubating 2 µg of the anti-FLAG M2 antibody (Sigma, F1804) with 20 µL of Dynabeads Protein G (Life Technologies) in 1x PBS containing 0.1% Triton X-100 and 0.5% Bovine Serum Albumin Fraction V (Sigma) for either 3-4 hours or overnight. After incubation, the antibody-attached beads were washed twice with Lysis300 buffer. After washing, 8 mg of WCL proteins were incubated with the anti-FLAG antibody-attached magnetic beads for 3-4 hours with gentle rotation. The beads then were washed four times with 200 µL of Lysis300 buffer. The proteins bound to the anti-FLAG antibody were eluted by incubating the beads with 100 µL of Lysis300 buffer containing 200 µg/mL of 3xFLAG peptide (Sigma) for one hour with gentle rotation. The sample then was placed in a magnetic stand, and the supernatant was moved to a new tube. These eluates were used as the IP-ORF2p samples.

Thirty µg of WCL or 3.5 µL of IP-ORF2p were separated using sodium dodecyl sulfate-polyacrylamide gel electrophoresis (SDS-PAGE) and transferred to a polyvinylidene difluoride (PVDF) membrane (Immobilon-LF PVDF [Merck Millipore]). The resultant membranes were incubated in Odyssey Blocking Buffer (LI-COR) and then incubated in 1x PBS containing 0.1% Tween-20 with the following primary antibodies: an anti-FLAG antibody (Sigma, F3165, 1:10,000 dilution); an anti-β-actin antibody (Santa Cruz Biotechnology, sc-47778, 1:10,000 dilution); and an anti-S6 ribosomal protein antibody (Cell Signaling Technology, 2217L, 1:5,000 dilution). After being washed four times with 1x PBS containing 0.1% Tween-20, the membranes were incubated in 1x PBS containing 0.1% Tween-20 and 0.01% SDS with the following secondary antibodies: IRDye 800CW Donkey anti-Mouse IgG (LI-COR, 925-32212, 1:10,000 dilution) and IRDye 680RD Donkey anti-Rabbit IgG (LI-COR, 925-68073, 1:10,000 dilution). Following incubation with the secondary antibodies, the membranes were washed four times with 1x PBS containing 0.1% Tween-20 and once with 1x PBS. Signals were detected using the Odyssey CLx imaging system (LI-COR) and analyzed with Image Studio Software (LI-COR).

### LEAP assays conducted with IP-ORF2p

An aliquot of IP-ORF2p was used in our original LEAP assays using a previously described protocol (58). Briefly, 2 µL of IP-ORF2p or Lysis300 buffer (negative control) was incubated with 200 µM dNTPs and 400 nM of the indicated oligonucleotides (**Table S1**: MN440, T4+VN-3’-OH; MN488, T4+AAddC; and MN503, T4+3T*invT) in 20 µL of LEAP buffer (50 mM Tris-HCl pH 8.0, 50 mM KCl, 5 mM MgCl_2_, 10 mM DTT, 0.05% Tween 20, 0.4 U/µL RNasin [Promega]) at 37°C for 60 minutes. The reaction was inactivated by heat treatment at 90°C for 10 minutes. Next, 1 µL of the resultant cDNA, or water (negative control), was amplified using AmpliTaq Gold 360 (Thermo Fisher Scientific) with the MN085 and MN485 oligonucleotides (**Table S1**) according to the protocol provided by the manufacturer using the following cycling conditions: one cycle: 95°C for 5 minutes; 30 cycles: 95°C for 15 seconds, 60°C for 30 seconds, and 72°C for 30 seconds; and a final extension cycle of 7 minutes at 72°C. The PCR products were separated on 2.5% agarose gels containing 1x TAE buffer and visualized with GelRed (Biotium). The LEAP products were excised from the gel using a razor blade, purified using a QIAquick Gel Extraction Kit (QIAGEN), and cloned into the pCR-II vector (Invitrogen). Plasmid DNAs were purified as described above and were subjected to Sanger DNA sequencing at either the University of Michigan DNA Sequencing Core facility or Eurofins Scientific.

### Production of recombinant L1 ORF2p-3xHA/3xFLAG baculovirus (Moran Laboratory)

The Bac-to-Bac Baculovirus Expression System (Invitrogen) was used to produce recombinant ORF2p-3xHA/3xFLAG using minor modifications to a previously described protocol (60). To obtain bacmid DNA, purified plasmid DNAs (pFastBac1 [Invitrogen], pMN001, pMN001_D205A, and pMN001_D702A) were transformed into DH10Bac cells (Invitrogen). The recombinant bacmid DNA was purified using a PureLink HiPure Plasmid DNA Miniprep Kit (Invitrogen) according to the protocol provided by the manufacturer. Positive clones were confirmed by PCR analysis and Sanger DNA sequencing at the University of Michigan DNA Sequencing Core facility. An aliquot (7 µL) of the purified recombinant bacmid was transfected into 850 µL *Sf*9 cells (at a concentration of 2 x 10^6^ cells/mL) using Cellfectin II (Invitrogen) according to the protocol provided by the manufacturer. The transfected cells then were incubated in either Insect-Xpress or Sf-900 II SFM medium supplemented with 10% FBS at 27°C, with shaking at 300 rpm, for one week. After centrifugation at 1,400 x *g* at 4°C for 15 minutes, the supernatant containing recombinant baculovirus (P1 virus stock) was transferred to a new tube and stored at 4°C in a tube covered with aluminum foil.

To amplify the recombinant baculoviruses, 100 µL of P1 virus stock was incubated with 50 mL of *Sf*9 cells (1 x 10^6^ cells/mL) at 27°C, with shaking at 150 rpm, for one week. The P2 virus stock was obtained by removing cells by centrifugation at 1,400 x *g* at 4°C for 15 minutes and collecting the supernatant. The titer of the virus stocks was determined using the baculoQUANT ALL-IN-ONE (OXFORD Expression Technologies).

### Production of recombinant L1 ORF2p-3xHA/3xFLAG and ORF2p-StrepII baculovirus (Barabas Laboratory)

The MultiBac System (Geneva Biotech) was used to produce recombinant ORF2p-3xHA/3xFLAG and ORF2p-StrepII proteins according to the protocol provided by the manufacturer with modifications. Briefly, to obtain bacmid DNA, purified plasmid DNAs (pMN001, p1253_ORF2p-StrepII, and p1393_ORF2p-D702A-StrepII) were transformed into DH10EmBacY cells (Geneva Biotech). The recombinant bacmid DNA was purified using a PureLink Quick Plasmid Miniprep Kit (Invitrogen) according to the protocol provided by the manufacturer up to the column binding step. DNA was precipitated with isopropanol to collect bacmid DNA. Ten or fifteen µg of the purified bacmid DNA was transfected into 3 mL *Sf*9 cells (at a concentration of 1.7-3.3 x 10^5^ cells/mL) using jetPRIME transfection reagent (Polyplus) according to the protocol provided by the manufacturer. The transfected cells then were incubated in Sf-900 III SFM medium (Gibco) at 27°C for 48-72 hours. When Yellow Fluorescent Protein (YFP) signal was visible in a few cells, the supernatant containing recombinant baculovirus (P1 virus stock) was transferred to a new tube and stored at 4°C covered with aluminum foil.

To amplify the recombinant baculoviruses, 3 mL of P1 virus stock was incubated with 25 mL of *Sf*9 cells (0.5 x 10^6^ cells/mL) at 27°C, with shaking at 120 rpm, for 48-72 hours until the cells showed YFP signals and the cell viability dropped to less than 85%. After centrifugation at 3,000 x *g* at 4°C for 10 minutes, the supernatant containing recombinant baculovirus (P2 virus stock) was transferred to a new tube, filtered with a syringe and a 0.22 µm pore size Millex syringe filter unit (Millipore), and stored at 4°C in a tube covered with aluminum foil. To further amplify the recombinant baculoviruses, 10 mL of P2 virus stock was incubated with 100 mL of *Sf*9 cells (1.0 x 10^6^ cells/mL) at 27°C, with shaking at 120 rpm, for 48-72 hours until the cells showed YFP signals and their viability dropped to less than 85%. After centrifugation at 3,000 x *g* at 4°C for 10 minutes, the supernatant containing recombinant baculovirus (P3 virus stock) was transferred to a new tube, filtered with a syringe and a 0.22 µm pore size Millex syringe filter unit (Millipore), and stored at 4°C in a tube covered with aluminum foil.

### Purification of rORF2p from High Five cells (Moran Laboratory)

The purification of rORF2p-3xHA/3xFLAG involved a three-step protocol using anti-FLAG affinity chromatography, anti-HA affinity chromatography, and heparin affinity chromatography. Each step of the rORF2p purification protocol was conducted on ice or at 4°C unless noted otherwise. Additional purification steps were added to remove detergents and nucleic acids that may interfere with functional rORF2p assays.

Briefly, either 250 mL or 333 mL of High Five cells (2 x 10^6^ cells/mL) were infected with recombinant rORF2p-3xHA/3xFLAG baculovirus at a multiplicity of infection (MOI) of 2.0 and incubated at 27°C, with shaking at 150 rpm, for 72 hours. The cells were harvested by centrifugation at 1,000 x *g* for 10 minutes at 4°C and stored at -80°C. The rORF2p-expressing High Five cells were thawed in 10 ml of Lysis Buffer (50 mM Tris-HCl pH8.0, 250 mM NaCl, 2.5 mM MgCl_2_, and 0.5% IGEPAL CA-630) containing 1x cOmplete EDTA-free Protease Inhibitor Cocktail, 1mM DTT, and 2.5 mL of Insect PopCulture reagent (Novagen). The lysed cells were resuspended using a pipette and then were sonicated at a 60% amplitude ten times at 20 second intervals with a 2-minute pause between each interval, using a Branson Sonifier cell disruptor 185 (Branson Ultrasonics). The sonicated cells were then incubated at room temperature for 30 minutes with gentle rotation in the presence of 8 U/mL benzonase (Novagen).

#### Anti-FLAG affinity chromatography

The sonicated cell solution was centrifuged at 12,000 x *g* for 30 minutes. The supernatant (10-12 mL) was collected and incubated with 400 µL of EZview Red ANTI-FLAG M2 Affinity Gel (Sigma) that was pre-equilibrated as a 50% volume/volume (V/V) affinity gel/Lysis buffer slurry for three hours with gentle rotation. The gel suspension was washed five times in 5 mL of Lysis Buffer (see above) by inverting the tubes. The solution was centrifuged at 1,400 x *g* after each wash, and the wash buffer was removed from the affinity gel using a pipette. For the elution of rORF2p, the anti-FLAG affinity gel was incubated with 800 µL of Lysis Buffer containing 250 µg/mL of a 3xFLAG peptide for one hour. The solution was centrifuged at 1,000 x *g* for one minute and the supernatant was transferred to a new tube and saved as a first eluate. The anti-FLAG affinity gel then was subjected to a second elution step by incubating the gel suspension with 500 µL of Lysis Buffer containing 250 µg/mL of FLAG peptide overnight with gentle rotation. As above, after centrifugation at 1,000 x *g* for one minute, the supernatant was transferred to the tube to combine with the first eluate.

#### Anti-HA affinity chromatography

Approximately 1.3 mL of the anti-FLAG affinity gel eluate was incubated with 200 µL of EZview Red ANTI-HA Affinity Gel (Sigma) for three hours with gentle rotation. The gel suspensions were washed five times; 1 mL of Lysis Buffer was mixed by inverting the tubes, the solution was centrifuged at 1,000 x *g*, and the buffer was removed from the affinity gel using a pipette. For the elution of rORF2p, the anti-HA affinity gel was incubated with either 500 or 600 µL of Lysis Buffer containing 250 µg/mL of an HA peptide (Sigma) for one hour. The solution was centrifuged at 11,000 x *g* for one minute and the supernatant was transferred to a new tube as a first eluant. The anti-HA affinity gel then was subjected to a second elution step by incubating the gel suspension with either 300 or 400 µL of Lysis Buffer containing 250 µg/mL of HA peptide overnight with gentle rotation. As above, after centrifugation at 11,000 x *g* for one minute, the supernatant was transferred to the new tube to combine with the first eluate. The combined eluates were cleared by centrifugation at 21,000 x *g* for 15 minutes.

#### Heparin affinity chromatography

Approximately 0.8 mL or 1 mL of the anti-FLAG/anti-HA affinity eluate was loaded onto a 1 mL HiTrap Heparin HP column (GE Healthcare) that was pre-equilibrated with Lysis Buffer. The column was then washed with five column volumes of Lysis Buffer. A six-column volume (∼6 mL) linear gradient of Lysis Buffer containing 250 mM to 1.5 M NaCl was applied to the column and was collected in 12 fractions of 500 µL. Three to four fractions (1.5-2.0 mL) containing eluted rORF2p (determined by Western blot, see below) were pooled and dialyzed against Lysis Buffer containing 250 mM NaCl in a Slide-A-Lyzer MINI Dialysis Device 20k MWCO (Thermo Fisher Scientific). The protein concentrations of the purified rORF2p solutions were determined using a reducing agent and detergent compatible colorimetric assay (*i.e.*, a BioRad RC DC Protein Assay). The solutions containing rORF2p, rORF2p_D205A, and rORF2p_D702A then were diluted with Lysis Buffer to either 0.5 ng/µL or 50 nM.

#### Additional removal of nucleic acids during purification of rORF2p by polyethyleneimine (PEI) precipitation

The rORF2p-expressing High Five cells were thawed in 10 ml of Lysis Buffer containing 750 mM NaCl, 1x cOmplete EDTA-free Protease Inhibitor Cocktail, 1mM DTT, and 2.5 mL of Insect PopCulture reagent and sonicated as described above. Benzonase treatment and centrifugation were performed as described above. PEI, at a final concentration of 0.04% w/v, was added to the supernatants and incubated for 10 minutes. Precipitates were removed by centrifugation at 12,000 x *g* for 30 minutes. The PEI precipitation protocol was repeated 3 to 4 times until the lysates were cleared of precipitates. The anti-FLAG affinity chromatography, anti-HA affinity chromatography, and heparin affinity chromatography were performed using the protocols outlined above with the modified buffers containing the following NaCl concentrations: 750 mM NaCl for incubation and wash steps conducted when using EZview Red ANTI-FLAG M2 Affinity Gel; 500 mM NaCl for elution with the FLAG peptide; and 500 mM NaCl for the incubation and wash steps conducted when using the EZview Red ANTI-HA Affinity Gel.

#### Further purification of rORF2p to remove IGEPAL CA-630

To remove IGEPAL CA-630 from purified rORF2p preps, which has the potential to inhibit ORF2p cDNA synthesis, the purification protocol without using PEI precipitation described above was repeated up to the elution step from the EZview Red ANTI-HA Affinity Gel chromatography with modified buffers containing the following NaCl concentrations: 750 mM NaCl for incubation and wash steps conducted when using EZview Red ANTI-FLAG M2 Affinity Gel; 500 mM NaCl for elution with the FLAG peptide; and 500 mM NaCl for the incubation and wash steps conducted when using the EZview Red ANTI-HA Affinity Gel. The supernatant from the anti-HA elution step was loaded onto a 1 mL HiTrap Heparin HP column that was pre-equilibrated in Lysis Buffer. The HiTrap Heparin HP column was washed with five column volumes of Lysis Buffer containing 250 mM NaCl followed by additional washes using twenty column volumes of Lysis Buffer containing 250 mM NaCl without IGEPAL CA-630. A six-column volume gradient of Lysis Buffer containing 250 mM to 1.5M NaCl without IGEPAL CA-630 was used to elute rORF2p; we collected 500 µL fractions as described above. The pooled samples containing rORF2p then were dialyzed against Lysis Buffer without IGEPAL CA-630 using a Slide-A-Lyzer MINI Dialysis Device 20k MWCO (Thermo Fisher Scientific). The protein concentration of the solution was determined using a BioRad RC DC Protein Assay. Glycerol was added to the purified rORF2p at a final concentration of 20% V/V followed by flash frozen in liquid nitrogen, and stored at -80°C.

#### Purification of rORF2p from High Five cells (Barabas Laboratory)

To determine the optimal infection condition, different volumes of the P3 virus (0.125-1 mL) were used to infect 10 mL of High Five cells (1.5 x 10^6^ cells/mL) followed by incubation at 27°C, with shaking at 120 rpm, for 5 days. The cell numbers, cell viability, and YFP signal were monitored daily. Thereafter, the recombinant baculovirus was used to infect 300 mL to 400 mL of High Five cells (1.5 x 10^6^ cells/mL) at the determined optimal ratio and incubated at 27°C, with shaking at 120 rpm, for the determined optimal time. Finally, the cells were pelleted by centrifugation at 1,000 x *g* for 10 minutes at 4°C. Most of the supernatant was removed but 30 mL were left to resuspend the cells. The resuspended cells were transferred to a new 50 mL tube and harvested by centrifugation at 1,000 x *g* for 10 minutes at 4°C. The supernatant was removed, the pellet was flash frozen in liquid nitrogen, and stored at -80°C.

Purification of ORF2p-StrepII and ORF2p-D702A-StrepII were performed on ice or at 4°C unless noted otherwise. The ORF2p-expressing High Five cells were thawed and lysed with 12.5 mL of Lysis Buffer 0.1% (50 mM Tris-HCl pH8.0, 250 mM NaCl, 2.5 mM MgCl_2_, 0.1% IGEPAL CA-630, 1mM DTT) containing 1x cOmplete EDTA-free Protease Inhibitor Cocktail, and 2.5 mL of Insect PopCulture reagent (Millipore). Cells were resuspended using a pipette and then were sonicated at a 40% amplitude (BRANSON Digital Sonifier 450) four times using the following cycling conditions: 30 cycles: 0.5 second ON and 0.5 second OFF, with a 2-to-3-minute pause between each cycle. The lysate then was incubated at room temperature for 30 minutes with gentle rotation in the presence of 8 U/mL benzonase (Millipore) to fragment genomic DNA and centrifuged in Beckman L-90K with rotor 45 TI at 125,000 x *g* for 45 minutes. The supernatant was collected, filtered using a 0.45 µm pore size Millex filter unit (Millipore) and loaded onto a 1 mL StrepTrap HP column (GE Healthcare) that was preequilibrated in Lysis Buffer 0.1%. The column was washed with 10 column volumes of Lysis Buffer 0.1%. The samples were eluted with 15 column volumes of Elution Buffer (50 mM Tris-HCl pH8.0, 250 mM NaCl, 2.5 mM MgCl_2_, 0.1% IGEPAL CA-630, 1mM DTT, 2.5 mM d-Desthiobiotin) and 1 mL fractions were collected from the column. SDS-PAGE was used to identify fractions containing ORF2p. Fractions containing ORF2p were pooled (∼4 mL) and loaded into 1 mL HiTrap Heparin HP column (GE Healthcare) pre-equilibrated in Lysis Buffer 0.1%. The column was washed with 20 column volumes of Wash Buffer (50 mM Tris-HCl pH8.0, 250 mM NaCl, 2.5 mM MgCl_2_), thereby removing the detergent from the samples. The sample was eluted with a six-column volume gradient from Wash Buffer to Heparin Elution Buffer (50 mM Tris-HCl pH8.0, 1.5 M NaCl, 2.5 mM MgCl_2_) and 1 mL fractions were collected. SDS-PAGE followed by Coomassie Brilliant Blue staining was used to identify fractions containing ORF2p. Fractions containing ORF2p were pooled (∼4 mL) and concentrated in a Vivaspin Turbo 4 10k MWCO concentrator (Sartorius) to 100 µL. The sample then was dialyzed using Slide-A-Lyzer MINI Dialysis Device 10k MWCO (Thermo Fisher Scientific) against 250 mL Wash Buffer. The protein concentration of purified ORF2p-StrepII and ORF2p-D702A-StrepII samples were determined by NanoDrop (Thermo Fisher Scientific) spectrophotometer. Samples were aliquoted, flash frozen in liquid nitrogen, and stored at -80°C.

Recombinant ORF2p-3xHA/3xFLAG was purified using the protocols outlined above with buffers containing 0.1% IGEPAL CA-630 and 250 mM NaCl up to the Anti-HA affinity gel chromatography step with the modifications noted below. The benzonase treated cells were centrifuged at 49,000 x *g* for 30 minutes and the supernatant was subjected to Anti-FLAG affinity chromatography. During Anti-FLAG purification, the samples were centrifuged at 3,000 x g in the wash step and at 8,000 x *g* in the elution step. During Anti-HA affinity purification, the samples were centrifuged at 8,000 x *g* in the wash and elution steps. The supernatant from the anti-HA elution step was loaded onto a 1-mL HiTrap Heparin HP column that was preequilibrated in Lysis Buffer 0.1%. The HiTrap Heparin HP column was washed with five column volumes of Lysis Buffer 0.1% followed by an additional wash using 20 column volumes of Wash Buffer. A six column volume gradient ranging from Wash Buffer to Heparin Elution Buffer was used to elute rORF2p and 1.0 mL fractions were collected. SDS-PAGE was used to identify fractions containing ORF2p. Fractions containing ORF2p were pooled (∼4 mL) and concentrated in a Vivaspin Turbo 4 10k MWCO concentrator (Sartorius) to 100 µL. The samples then were dialyzed against 250 mL Wash Buffer in a Slide-A-Lyzer MINI Dialysis Device 10k MWCO (Thermo Fisher Scientific). The protein concentration of the solution was determined using a NanoDrop (Thermo Fisher Scientific) spectrophotometer. Glycerol was added to the purified rORF2p at a final concentration of 20% (V/V). The sample was aliquoted, flash frozen in liquid nitrogen, and stored at -80°C.

### Visualization of rORF2p

Either 4 ng or 200 fmol of purified rORF2p was separated using SDS-PAGE and visualized using the SilverQuest Staining Kit (Invitrogen) according to the protocol provided by the manufacturer. Western blotting was used to further confirm the presence of rORF2p. Briefly, either 6 ng or 100 pg of purified rORF2p were separated by SDS-PAGE and transferred to Immobilon-LF PVDF membranes. The membranes were incubated in Odyssey Blocking Buffer and then incubated in 1x PBS containing 0.1% weight per volume (w/v) Tween-20 with the following primary antibodies: an anti-FLAG antibody (1:10,000 dilution), an anti-HA antibody (Roche, 11867423001, 1:3,000 dilution), and an anti-hORF2N antibody (Moran Laboratory; directed against ORF2p amino acids 154-167; DRSTRQKVNKDTQE, 1:3,000 dilution). After being washed four times with 1x PBS containing 0.1% Tween-20, the membranes were incubated in 1x PBS containing 0.1% Tween-20 and 0.01% SDS with the following secondary antibodies: IRDye 800CW Donkey anti-Mouse IgG (1:10,000 dilution), IRDye 800CW Donkey anti-Rabbit IgG (LI-COR, 925-32213, 1:10,000, dilution), and IRDye 800CW Goat anti-Rat IgG (LI-COR, 926-32219, 1:10,000 dilution). After incubation with secondary antibodies, the membranes were washed four times with 1x PBS containing 0.1% Tween-20 and once with 1x PBS. Signals were detected using the Odyssey CLx imaging system and analyzed with Image Studio Software.

### *In vitro* synthesis of RNA templates for the modified LEAP reaction

To synthesize DNA templates to generate RNAs used in modified LEAP reactions, we used PCR using the following oligonucleotide primer and DNA plasmid combinations (**Table S1**: MN056, MN058, and pJCC5/L1.3 to amplify the 3U RNA template; MN056, MN057, and pJCC5/L1.3 to amplify the 3U+23A and 3U+polyA RNA templates; MN470, MN468, and pJJ101/L1.3 to amplify the JS RNA template; MN470, MN469, and pJJ101/L1.3 to amplify the JS+23A and JS+polyA RNA templates). PCR reactions were conducted using either Platinum Pfx DNA Polymerase or Platinum SuperFi DNA Polymerase (Invitrogen). The PCR-amplified templates were purified from agarose gels using a QIAquick Gel Extraction Kit.

For *in vitro* transcription reactions, approximately 1 µg of template DNA was used in a 30 μL reaction using either a T7 Quick High Yield RNA Synthesis Kit or a HiScribe T7 Quick High Yield RNA Synthesis Kit (New England BioLabs). The reaction mixtures were incubated at 37°C for 16 hours and then treated with 4 units of DNase I at 37°C for 30 minutes to remove the DNA template. The resultant RNAs were gel-purified from 7% polyacrylamide/urea (8M) denaturing gels in 1x TBE buffer. The region of the gel containing RNAs was excised using a razor blade. The gel slice then was incubated with 1 mL of TE (10 mM Tris-HCl, 1 mM EDTA, pH 8.0) at 4°C with gentle rotation overnight to elute the RNAs. The eluted RNAs were purified using ethanol precipitation. For the 3U+polyA and JS+polyA RNAs, poly(A) tails were added using *E. coli* Poly(A) Polymerase after gel purification (New England BioLabs). The polyadenylated RNAs then were purified using phenol chloroform extraction and ethanol precipitation. To verify transcript integrity, 200 ng of RNA was separated on 7% polyacrylamide/urea (8M) denaturing gel and visualized using GelRed.

### Modified LEAP reactions using purified rORF2p

An aliquot (2 µL) of purified rORF2p was incubated with 500 nM of the indicated oligonucleotide (**Table S1**: MN440, T4+VN-3’-OH; MN488, T4+AAddC; MN503, T4+3T*invT; MN746, T4+3T*ddC), 1 ng of the indicated *in vitro* transcribed RNA, and 200 μM dNTPs, in 10 µL of LEAP buffer at 37°C for 60 minutes. The reaction was inactivated by heat treatment at 90°C for 20 minutes. An aliquot (1 µL) of the resultant cDNA or water (negative control) was PCR amplified using the following oligonucleotides (**Table S1**: ORF2M and MN085 for cDNAs generated using the 3U, 3U+23A and 3U+polyA RNA templates; MN462 and MN085 for cDNAs generated using the JS, JS+23A, and JS+polyA RNA templates).

To amplify the cDNA products made by rORF2p derived from the MN440, MN488, and MN503 oligonucleotides (**Table S1**) in modified LEAP assays, we used Universe High-Fidelity Hot Start DNA Polymerase (Bimake) according to the protocol provided by the manufacturer using the following cycling conditions: one cycle: 95°C for 3 minutes; 25 cycles: 95°C for 15 seconds, 58°C for 15 seconds, and 72°C for 20 seconds; and a final extension cycle of 5 minutes at 72°C. To amplify the cDNA products made by other versions of rORF2p (*i.e.*, rORF2p+IGEPAL-PEI, rORF2p+IGEPAL+PEI, and rORF2p-IGEPAL-PEI) from the MN 746 oligonucleotide (**Table S1**), we used AmpliTaq Gold 360 according to the protocol provided by the manufacturer using the following cycling conditions: one cycle: 95°C for 5 minutes; 30 cycles: 95°C for 15 seconds, 58°C for 30 seconds, and 72°C for 30 seconds; and a final extension cycle for 7 minutes. We also amplified the cDNA products derived from the primers mentioned above with Platinum SuperFi DNA Polymerase according to the protocol provided by the manufacturer using the following cycling conditions: one cycle: 98°C for 2 minutes; 30 cycles: 98°C for 10 seconds, 58°C for 10 seconds, and 72°C for 30 seconds; and a final extension cycle for 5 minutes.

The cDNA products from the modified LEAP reactions were separated on 2.5% agarose gels containing 1x TAE buffer and visualized with GelRed. The products were excised from the gels using a razor blade, purified using a QIAquick Gel Extraction Kit, and cloned into the pCR-Blunt, pCR-II, pCR4-TOPO, or pCR4Blunt-TOPO vectors (Invitrogen). Plasmid DNAs were purified as noted above and subjected to Sanger DNA sequencing at the University of Michigan DNA Sequencing Core facility or Eurofins Scientific. Importantly, the PCR reactions conducted with AmpliTaq Gold 360 and Platinum SuperFi DNA Polymerase gave similar results.

### Modified LEAP reactions conducted with MMLV RT and *in vitro* synthesized RNA

The modified LEAP reaction conducted with MMLV RT (Promega) was performed according to the protocol provided by the manufacturer. Briefly, 8 units of MMLV RT or water (negative control) was incubated with 500 nM of the indicated ssDNA oligonucleotide (**Table S1**: MN440, T4+VN; MN488, T4+AAddC; MN503, T4+3T*invT; MN746, T4+3T*ddC) and 1 ng of the indicated RNA in 10 µL 1x Reaction Buffer at 37°C for 60 minutes. The resultant cDNAs were amplified and analyzed as described in the “Modified LEAP reactions using purified rORF2p” section above.

### Purification of L1 RNPs for use in LEAP assay

The L1 RNP purification was modified slightly from previously described protocols (41,61). Briefly, HeLa-JVM cells were plated in a T175 flask containing DMEM-complete medium at approximately 6 x 10^6^ cells per flask. The following day (day 0), the cells were transfected with 30 µg of the indicated plasmids using 90 µL of the FuGENE6 transfection reagent and 3 mL of Opti-MEM according to the protocol provided by the manufacturer. The following day (day 1), the transfection medium was replaced with fresh DMEM-complete medium. On days 3-9, the medium was replaced daily with DMEM-complete medium containing 200 µg/mL hygromycin B. On day 9, the cells were collected, divided between two tubes, washed with cold 1x PBS, and flash-frozen in a dry ice/ethanol bath.

The following steps were performed on ice or at 4°C unless noted otherwise. A 1 mL solution of Lysis RNP buffer (5 mM Tris-HCl pH 7.5, 2.5 mM MgCl_2_, 1.5 mM KCl, 1% deoxycholic acid, 1% Triton X-100, 1x cOmplete EDTA-free Protease Inhibitor Cocktail) was added to the frozen cell pellets and the cells were incubated for 15 minutes. To obtain WCLs, the insoluble cell debris was removed by centrifugation at 1,400 x *g* for 10 minutes. The WCLs then were subjected to ultracentrifugation though a sucrose cushion (8.5% and 17% sucrose) at 168,000 x *g* for two hours. The RNP pellets were resuspended in 1x cOmplete EDTA-free Protease Inhibitor Cocktail, and the protein concentration then was determined using the Bradford reagent assay. The L1 RNPs were diluted to 1 µg/µL in 1x cOmplete EDTA-free Protease Inhibitor Cocktail in a 50% glycerol V/V solution and were stored at -20°C.

An aliquot of WCLs (30 µg) or L1 RNPs (3.5 µg) then were subjected to SDS-PAGE and western blot analysis as described above. The following primary antibodies were used: ɑ-N-ORF1p antibody (made in the Moran Laboratory; directed against ORF1p amino acids 31-49; EQSWMENDFDELREEGFRR, 1:5,000 dilution); an anti-S6 ribosomal protein antibody (1:5,000 dilution); and an anti-β-actin antibody (1:10,000 dilution). The following secondary antibodies were used: IRDye 800CW Donkey anti-Rabbit IgG (1:10,000 dilution) and IRDye 680RD Goat anti-Mouse IgG (LI-COR, 926-68070, 1:10,000 dilution).

### LEAP assays conducted with L1 RNPs

The original version of the LEAP assay used in this study was previously described in the following publications (41,54,56). Briefly, a 1 µg aliquot of an L1 RNP sample, the indicated oligonucleotide primer, and dNTPs were incubated in LEAP buffer at 37°C for one hour. The resultant cDNAs then were amplified using PCR, separated using a 2% agarose gel, and visualized by staining with ethidium bromide as described in previous publications (41,54,56).

### RT-PCR reactions to detect L1 RNAs in L1 RNP samples

RNA was isolated from L1-RNPs derived from HEK293T cells transfected with pJJ101/L1.3 using a RNeasy Mini Kit (QIAGEN). The L1 RNA was reverse transcribed using MMLV RT (Promega) or AMV RT (Promega) and the indicated oligonucleotides as described in previous publications (41,54). The resultant cDNAs were amplified by PCR, separated using a 2% agarose gel, and visualized by staining with ethidium bromide as described above.

### Exonuclease treatment of [γ^32^-P]-ATP labeled oligonucleotides

Oligonucleotides (**Table S1**: MN440, T4+VN-3’-OH; MN488, T4+AAddC; and MN503, T4+3T*invT) were radiolabeled using [γ^32^-P]-ATP (Perkin Elmer [now named Revvity]) and T4 Polynucleotide Kinase (New England BioLabs) according to the protocol provided by the manufacturer. Free [γ-^32^P]-ATP was removed from the solution using a MicroSpin G-25 Column (Cytiva) according to the protocol provided by the manufacturer. Approximately 200 fmol (0.5 µL) of each oligonucleotide was incubated in the presence (+) or absence (-) of 10 U of Exonuclease I (New England BioLabs) in 5.5 µL of LEAP buffer at 37°C for the indicated time. The reaction solutions then were incubated with 5.5 µL of 2x TBE-Urea Sample Buffer (BIO-RAD) at 75°C for 10 minutes and separated using a 12% polyacrylamide/8M Urea denaturing gel. The gel was rinsed in distilled water for 30 minutes and then dried on Whatman paper under vacuum using Slab Gel Dryer SGD5040 (Savant) without heat. Autoradiography allowed the visualization of the γ^32^-P signal.

### RT-independent ligation-based assay to monitor alt-EN activity

#### The oligonucleotide 3’ end processing reaction

ORF2 protein aliquots (1.5 µL of an L1 RNP, 1.5 µL of IP-ORF2p, or 2 µL of rORF2p) were incubated with 1 µM of the indicated oligonucleotide (**Table S1:** MN528, T4+AAddC; MN527, T4+3T*invT; MN526, T4-3’-OH) in 15 µL of LEAP buffer at 4°C for 15 minutes and then at 37°C for 60 minutes. The reaction was inactivated by heat treatment at 90°C for 20 minutes. RNAs were removed by incubating the sample in a solution containing 20 µg of RNase A (DNase and protease-free, Thermo Fisher Scientific), 20 U of RNase I (Promega), and 1x RNase I buffer at 37°C for two hours. The RNases then were inactivated by heat treatment at 70°C for 20 minutes.

#### Purification of the oligonucleotide

Oligonucleotides from the above reactions were purified using a Dynabeads kilobaseBINDER Kit (Invitrogen). Briefly, 22 µL of the oligonucleotide containing sample was incubated with 20 µL of a Dynabeads M-280 Streptavidin solution (Invitrogen), which was first pre-equilibrated using the Binding Solution in the kit, for three hours with gentle shaking at room temperature. The beads were washed twice with 30 µL of Washing Solution (10 mM Tris-HCl pH 7.5, 1 mM EDTA, and 2 M NaCl), once with 30 µL of water, twice with 30 µL of 0.15 M NaOH for two minutes, and twice with 30 µL of TE buffer. The purified bead solutions containing the bound DNA oligonucleotides then were incubated at 95°C for 10 minutes in 30 µL of TE.

#### Additional samples

For pre-incubation steps with or without RNase, the pJJ101/L1.3 RNP and pTMO2F3 IP-ORF2p samples were incubated in the presence or absence of 7.5 µg of RNase A and 0.75U of RNase I at 37°C for 15 minutes prior to the 3’ end processing reaction. For heat-inactivated samples (pJJ101/L1.3_hi, pTMO2F3 _hi, and rORF2p_hi), the pJJ101/L1.3 RNP, pTMO2F3 IP-ORF2p, and WT rORF2p were incubated at 90°C for 20 minutes prior to conducting the 3’ end processing reaction. For 3’ end processing reactions conducted with rORF2p in the presence or absence of exogenous RNA, rORF2p was pre-incubated in the presence of 23A RNA (**Table S1**: MN584) or 23U RNA (**Table S1**: MN585), or without exogenously added RNA, at room temperature for 15 minutes.

#### The ligation reaction

An aliquot (3.6 µl) of the purified oligonucleotide from the 3’ end processing reaction was incubated with an oligonucleotide (1.0 pmol) containing a 5’-phosphate (**Table S1**: MN544) and a “splint” or “bridging” oligonucleotide (**Table S1**: MN546) at 65°C for 10 minutes, and 25°C for 3 minutes, to facilitate annealing of the oligonucleotides in the mixture. The annealed oligonucleotide mixture then was incubated either with or without 0.2 µL of HiFi Taq DNA ligase (New England BioLabs) in 20 µL of 1x HiFi Taq DNA Ligase Buffer. The ligation reaction was conducted in a thermocycler using the following cycling conditions: 15 cycles: 37°C for 15 minutes followed by 60°C for one minute. The ligated DNA was purified using a Dynabeads kilobaseBINDER Kit. Briefly, 20 µL of the oligonucleotide containing sample was incubated with 5 µL of a Dynabeads M-280 Streptavidin solution (Invitrogen), which was first preequilibrated using the Binding Solution in the kit, for three hours with gentle shaking at room temperature. The beads were washed twice with 40 µL of Washing Solution, once with 40 µL of water, twice with 40 µL of 0.15 M NaOH for two minutes, and twice with 40 µL of 1xTE buffer. The purified bead solution containing the bound DNA oligonucleotide then was incubated at 95°C for 10 minutes in 40 µL of 1xTE buffer.

#### Taqman PCR to detect the ligated product

An aliquot (1 µL) of the purified bead solution containing ligated DNA was subjected to TaqMan qPCR. The qPCR reactions were conducted using an ABI 7300 Real-Time PCR System (Thermo Fisher Scientific). Each reaction contained 5 µM of oligonucleotide PCR primers (**Table S1**: MN552 and MN553) and 2.5 µM of the Taqman probe (**Table S1**: MN551) in a TaqMan Universal PCR Master Mix (Applied Biosystems). PCR was conducted using the following cycling conditions: one cycle: 50°C for 2 minutes followed by 95°C for 10 minutes; 40 cycles: 95°C for 15 seconds and 60°C for 1 minute. Each experiment was performed with three technical replicates at least three independent times. A standard curve was generated using serial 10-fold dilutions of the pT4Zf1 plasmid. In each experiment, the relative amount of ligated DNA was normalized to the buffer control. A Student’s t-test was used to determine the reported p-values.

As control experiments, we used the above protocol but removed one component from the reaction mixture (*i.e.*, the “splint [bridge or junction]” oligonucleotide [**Table S1**: MN546], the 5’ phosphorylated oligonucleotide [**Table S1**: MN544], or the substrate oligonucleotide [*e.g.*, **Table S1**: MN526]). The removal of any of the listed components decreased the amount of the ligated DNA product to background levels.

### Modified LEAP reactions conducted with different L1 ORF2p preparations

The indicated L1 ORF2p preparations in the following buffers were used in these experiments: FLORF2p: 25mM HEPES pH 8, 500mM KCl, 5% glycerol, 2mM MgOAc, 0.5mM TCEP (tris [2-carboxyethyl] phosphine hydrochloride); and ORF2pCore: 20 mM HEPES pH 8.0, 500 mM NaCl, 5% glycerol, 1 mM DTT, 2 mM MgCl_2_. Both proteins were generous gifts from Dr. Martin S. Taylor and Dr. Trevor van Euwen (26).

An aliquot of FLORF2p or the ORF2pCore protein (10 nM or indicated concentration) was incubated with 500 nM of the indicated oligonucleotide (**Table S1**: MN440, T4+VN-3’-OH; MN488, T4+AAddC; MN503, T4+3T*invT; or MN746, T4+3T*ddC), 1 ng of the indicated RNA template, and 200 µM dNTPs in 10 µL of LEAP buffer at 37°C for 60 minutes. The reaction was inactivated using heat treatment at 90°C for 20 minutes. A water (no protein) sample served as a negative control.

An aliquot (1 µL) of the resultant cDNA was PCR amplified using AmpliTaq Gold 360 and the MN462 and MN085 oligonucleotides (**Table S1**) according to the protocol recommended by the manufacturer using the following cycling conditions: one cycle: 95°C for 5 minutes; 30 cycles 95°C for 15 seconds, 58°C for 30 seconds, and 72°C for 30 seconds; and a final extension cycle of 7 minutes at 72°C. We also amplified the above cDNA products using Platinum SuperFi DNA Polymerase according to the protocol provided by the manufacturer using the following cycling conditions: one cycle: 98°C for 2 minutes; 30 cycles: 98°C for 10 seconds, 58°C for 10 seconds, and 72°C for 30 seconds; and a final extension cycle for 5 minutes.

The products from the modified LEAP reactions were separated on 2.5% agarose gels containing 1x TAE buffer and visualized with GelRed. The LEAP products were excised from the gels using a razor blade, purified using a QIAquick Gel Extraction Kit, and cloned into the pCR4-TOPO or pCR4Blunt-TOPO vector (Invitrogen). Plasmid DNAs were prepared as described above and were subjected to Sanger DNA sequencing at the University of Michigan DNA Sequencing Core facility or Eurofins Scientific. Characterization of the cDNA products obtained from PCR reactions using AmpliTaq Gold 360 and Platinum SuperFi DNA Polymerase yielded similar results.

### Modified LEAP reactions conducted with Induro RT

One unit of Induro RT (New England BioLabs), one unit of Ultrapure SMART MMLV RT (TaKaRa], or 10 nM of rORF2p was incubated with 500 nM of the indicated oligonucleotide (**Table S1**: MN440, T4+VN-3’-OH; MN488, T4+AAddC; MN503, T4+3T*invT; or MN746, T4+3T*ddC), 1 ng of the JS+23A RNA, and 200 µM dNTPs in 10 µL of LEAP buffer at 37°C, or the otherwise indicated temperature, for 60 minutes. The reaction was inactivated using heat treatment at 90°C for 20 minutes. Heat-inactivated proteins were incubated at 90°C or 95°C for 20 minutes before conducting the modified LEAP reaction. Proteinase K-treated proteins were incubated with 80 U/mL of Proteinase K (New England BioLabs) at 37°C for 30 minutes, and then at 95°C for 10 minutes, before conducting the modified LEAP reaction. A water (no protein sample) served as a negative control. An aliquot (1 µL) of the resultant cDNA was amplified using AmpliTaq Gold 360 or Platinum SuperFi DNA Polymerase and the MN462 and MN085 oligonucleotides (**Table S1**) as noted in the above section. Characterization of the cDNA products obtained from PCR reactions using AmpliTaq Gold 360 and Platinum SuperFi DNA Polymerase yielded similar results.

### Complementation reverse transcription-PCR (RT-PCR) assay

MMLV RT (RNase H-) (Promega M5301) and Induro RT (NEB M0681S) were diluted to 10 U/µL in RT Buffer (50 mM Tris-HCl pH 7.5, 50 mM KCl, 5 mM MgCl_2_, and 0.05% Tween-20). ORF2p-StrepII and ORF2p-D702A-StrepII (RT [-]) were diluted to 45 ng/µL in RT Buffer. For complementation reactions, 1 µL of ORF2p-StrepII, ORF2p-D702A-StrepII or Induro RT were mixed with 1 µL of MMLV RT in PCR tubes. For single enzyme reactions, 2 µL of each enzyme was pipetted in PCR tubes. Aliquots (2 µL) of RT Buffer were used for negative controls. All enzymes and controls were incubated with 333 nM of the indicated ssDNA oligonucleotide (**Table S1**: MN503, T4+3T*invT; 4228, T6-3’-OH), 100 ng of JS+23A RNA, 166 µM of dNTPs, 8.3 mM of DTT, and 1.6 U of RiboLock (Thermo Fisher Scientific EO0381) in RT Buffer in 12 µL total volume (2 µL enzymes + 10 µL reaction components) at 37°C for 60 minutes. The reactions were inactivated by heat treatment at 90°C for 20 minutes. Then, a 1 µL aliquot of each heat inactivated reaction was used for PCR amplification of cDNA with the oligonucleotide primers (**Table S1**: MN462 and MN085) and KOD One™ PCR Master Mix (Toyobo) according to the protocol provided by the manufacturer. The following cycling conditions were used: one cycle: 98°C for 20 seconds; 25 cycles: 98°C for 10 seconds, 60°C for 5 seconds, and 68°C for 1 second; a final extension period of 5 seconds at 68°C. The final PCR products were separated by electrophoresis on 2% agarose gels with 1x TAE buffer and visualized by staining with ethidium bromide.

## Results

### Immunoprecipitated L1 ORF2p is associated with an alternative nuclease activity

We previously observed that L1 RNPs isolated by ultracentrifugation through a sucrose cushion have a nuclease activity that could remove a 3’ modification that blocks cDNA synthesis from an oligonucleotide, before it is used to prime the reverse transcription of L1 mRNA in a LEAP reaction (54).

However, because these RNP preparations also contained host proteins and host RNAs, we could not determine whether this nuclease activity was associated with ORF2p.

To explore whether the nuclease activity was contained within ORF2p, we first isolated ORF2p by immunoprecipitation from whole cell lysates derived from human embryonic kidney (HEK293T) cells transfected with a monocistronic WT ORF2p expression construct or monocistronic mutant constructs that harbor missense mutations in the L1 ORF2p EN or RT domains (**Fig. 1A**). To specifically precipitate ORF2p, we exploited an epitope tag (3xFLAG), which is compatible with retrotransposition of a full-length L1 when present at the carboxyl-terminus of ORF2p (ORF2p-3xFLAG) (58). The resultant immunoprecipitated ORF2p samples appeared to be considerably cleaner than crude RNP preparations (**Fig. S1**) and were examined for their ability to promote L1 mRNA cDNA synthesis using LEAP assays conducted with different oligonucleotide primers (**Fig. 1B and Fig. S1A**).

**Fig. 1:**
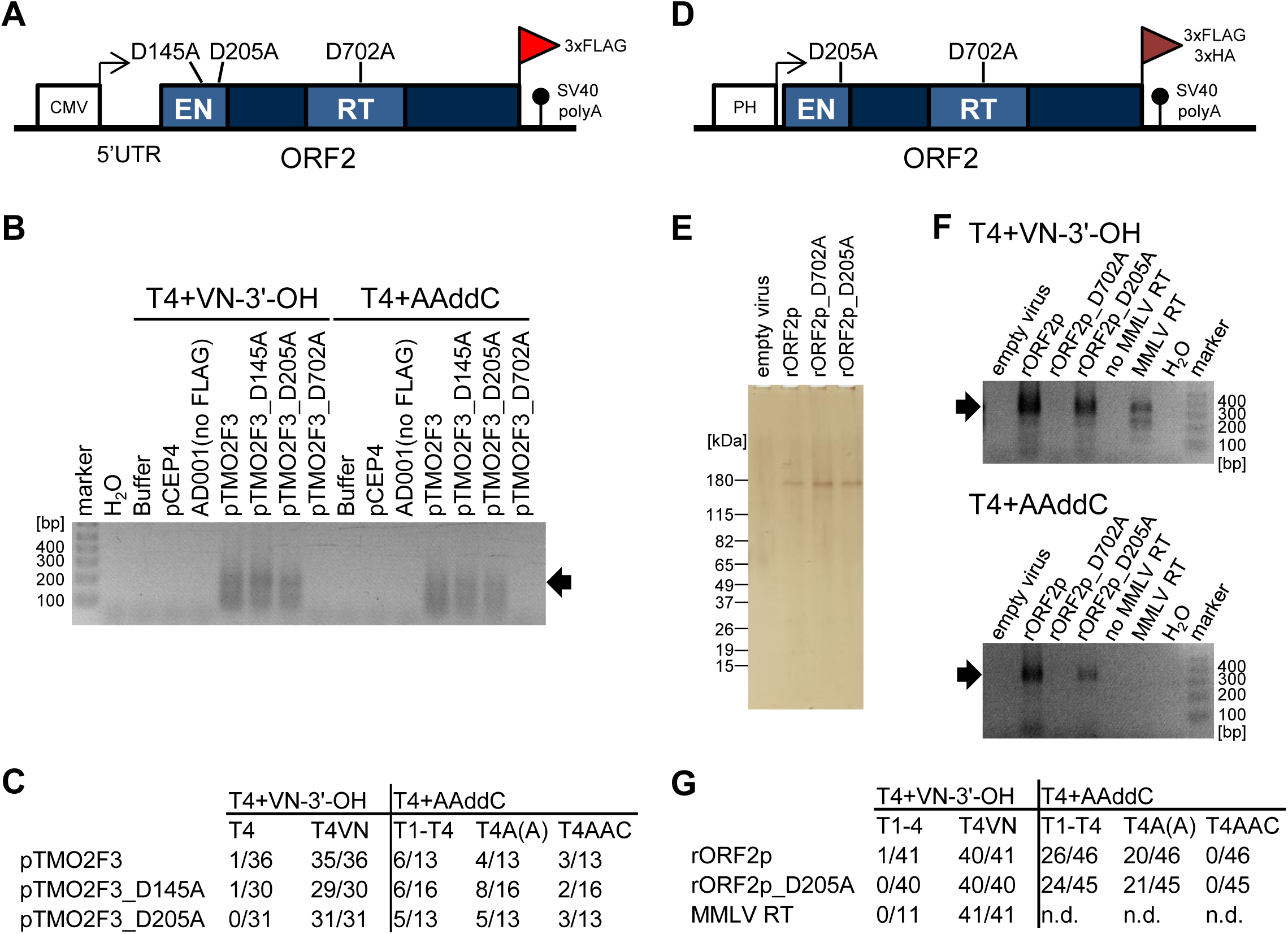
Immunoprecipitated (IP-ORF2p) and recombinant (rORF2p) contain a nuclease activity that can process a 3’ end modification from an oligonucleotide in a LEAP or modified LEAP reaction. ***(A)*** *Schematic* of the monocistronic L1.3 ORF2p expression construct. The L1.3 5’UTR (black line), *L1.3 ORF2* (blue rectangle) containing an in-frame 3x-FLAG epitope tag at its 3’ end (red flag), and 3’UTR (black line) were cloned into the pCEP4 expression vector, creating pTMO2F3. The pCEP4 vector contains a CMV immediate early promoter (white rectangle and black arrow) and an SV40 polyadenylation signal (black lollipop), which facilitates L1 expression. The approximate positions of missense mutations in the endonuclease (EN) and the reverse transcriptase (RT) domains (white lettering in light blue rectangles) are indicated in the schematic. ***(B)*** *Results from the LEAP assay*. Immunoprecipitated ORF2p (IP-ORF2p) was used in LEAP assays containing the following oligonucleotides with different 3’ ends: T4+VN-3’-OH (MN440) and T4+AAddC (MN488) (see **Table S1** for complete sequences). The L1 transfected into human HEK293T cells is indicated above the agarose gel image. The black arrow indicates the expected size of the full-length LEAP product. Note: we previously observed and characterized smaller cDNA products in LEAP reactions conducted in the absence of L1.3 ORF1p (41,54). Negative controls: water (H_2_O, PCR control), buffer alone (LEAP control), and the pCEP4 empty vector (LEAP control). Marker, molecular weight size controls in base pairs (bp). At least three independent biological replicates were conducted for each transfection condition. ***(C)*** *Characterization of LEAP products.* The number of products sequenced from LEAP reactions in panel B using the T4+VN-3’-OH and T4+AAddC oligonucleotides are indicated in the Table (denominator). LEAP products exhibiting a 3’-end modification (numerator) and the inferred 3’-end used to prime L1 cDNA synthesis is indicated at the top of each column. Note: some LEAP products from reactions conducted with the T4+AAddC oligonucleotide contained a cytosine residue (T4AAC) at the oligonucleotide/cDNA junction, which likely resulted from the reverse transcription of a guanosine ribonucleoside present in the L1 RNA template. ***(D)*** *A schematic of the recombinant ORF2p (rORF2p) expression construct.* Transcription is augmented by a polyhedrin promoter (PH, black arrow) and SV40 poly(A) signal. *L1.3 ORF2* (see panel A for labeling details) contains an in-frame 3x-FLAG/3x-HA tag at its the 3’ end (maroon flag). **(*E)*** *Purification of rORF2p from insect cells*. Recombinant ORF2p was expressed in High Five cells using the Bac-to-Bac Baculovirus Expression System. The resultant WT and mutant rORF2 proteins were purified using: (**i**) anti-FLAG affinity chromatography, (**ii**) anti-HA affinity chromatography, and (**iii**) heparin column chromatography (see **Materials and Methods**). The rORF2p proteins were separated on a denaturing polyacrylamide gel. Silver staining was used to visualize the proteins. Negative control, empty virus. Molecular weight controls in kilodaltons (kDa) are indicated at the left of the gel image. ***(F)*** *Recombinant WT and EN-mutant, but not RT-mutant, proteins have nuclease activity.* Modified LEAP reactions (**see Fig. S2D**) were conducted with WT and mutant rORF2p using the T4+VN-3’-OH (top) or T4+AAddC (bottom) oligonucleotide primer and JS+polyA RNA template. The labeling is like that described in panel B. The black arrow indicates the expected size of the full-length LEAP product. Negative controls: empty virus (modified LEAP assay control), no MMLV RT (modified LEAP assay control), and water (H_2_O, PCR control). Note: MMLV could only use the T4+VN-3’-OH oligonucleotide to prime cDNA synthesis. The rORF2p D702A mutant lacked activity in the modified LEAP reactions. Marker, molecular weight size controls (bp). At least three independent biological replicates were conducted for each reaction condition. ***(G)*** *Characterization of Products from the modified LEAP reaction*. The labeling is like that described in Fig.1C.

LEAP activity was readily detected using the immunoprecipitated WT (pTMO2F3) and L1 EN-mutant (pTMO2F3_D145A or pTMO2F3_D205A) samples, but not in L1 RT-mutant (pTMO2F3_D702A), when we used a DNA oligonucleotide primer that ends in a 3’-OH (**Fig. 1B** and **Table S1**: T4+VN-3’-OH, where V = A, C, or G and N = A, C, G, or T) or a dideoxynucleoside (**Fig. 1B** and **Table S1**: T4+AAddC, where ddC indicates a 3’-dideoxycytosine). Characterization of the LEAP products from reactions using the T4+VN-3’-OH primer revealed that L1 cDNA synthesis began within either the 3’ L1 poly(A) tail or the L1 3’UTR; 95/97 LEAP products initiated cDNA synthesis at a “VN” nucleotide pair (**Fig. 1C** and **Supplemental Dataset**). The two other products appeared to be processed by a nuclease prior to being used to prime L1 cDNA synthesis (**Fig. 1C** and **Supplemental Dataset**), which is consistent with our previous results (41).

Characterization of the LEAP products from reactions using the T4+AAddC primer revealed that L1 cDNA synthesis again initiated within either the 3’ L1 poly(A) tail or the L1 3’UTR. Notably, examination of the primer/L1 junction sequences revealed that 34/42 LEAP products exhibited evidence of processing at their 3’ end prior to being used to prime L1 cDNA synthesis (**Fig. 1C**). A small fraction of products (**Fig. 1C**, last column and **Supplemental Dataset**) contained a cytosine nucleoside at the primer/cDNA junction. However, we could not distinguish whether the cytosine was derived from the T4+AAddC primer or was part of the cDNA product synthesized from the L1 RNA template, which contains a complementary guanosine at the relevant position. As expected, the immunoprecipitated ORF2p L1 RT-mutant (D702A) lacked activity in the LEAP assay. Thus, consistent with our previous observations using crude RNP preparations, both the WT and L1 EN-mutant immunoprecipitated preparations appear to contain a nuclease activity that can remove a mismatch or ddC from an oligonucleotide prior to its use as a primer in a LEAP reaction (54).

### WT and EN mutant recombinant ORF2p are associated with a nuclease activity

To further explore whether the alternative nuclease activity was contained within ORF2p or is associated with a host cellular protein, we purified full-length recombinant WT, EN-mutant (D205A), and RT-mutant (D702A) proteins containing a carboxyl-terminal 3xFLAG/3xHA epitope tag from insect cells (**Fig. 1D** and **Fig. 1E**, and **Fig. S2A**, **Fig. S2B,** see **Methods**). We then incubated an aliquot of a purified rORF2p sample, the T4+VN-3’-OH oligonucleotide (**Table S1**), and an *in vitro* transcribed RNA derived from the endogenous (3U) and engineered (JS) L1 3’UTR that either contained (**Fig. S2C**, 3U + 23A, 3U + polyA, JS+23A, or JS+polyA) or lacked (**Fig. S2C**, 3U or JS) a 3’ poly(A) tail (**Fig. S2C**) in LEAP buffer containing dNTPs (**Fig. S2D**, herein referred to as the modified LEAP assay).

WT and EN-mutant rORF2p (rORF2p_D205A), but not the RT-mutant (rORF2p_D702A), could efficiently use the T4+VN-3’-OH oligonucleotide to prime the synthesis of L1 cDNAs from RNA templates ending in a 3’ poly(A) tail (**Fig. 1F**, JS+polyA, **Fig. S2E**, 3U+23A or 3U+polyA, and **Fig. S2F**, JS+23A) or from RNA templates lacking a poly(A) tail (**Fig. S2E** and **Fig. S2F**, 3U and JS, respectively), albeit at lower levels, in modified LEAP reactions. Sequencing revealed that 40/41 of the WT rORF2p and 40/40 EN-mutant (rORF2p_D205A) modified LEAP products derived from the RNA templates initiated cDNA synthesis at a “VN” nucleotide pair (**Fig. 1G** and **Supplemental Dataset**). Moreover, sequencing revealed that the cDNAs derived from the 3U and JS RNA templates lacking a poly(A) tail are due to alternative ORF2p priming sites at short regions of complementarity present at the primer/template junction near the 3’ end of those RNAs (**Fig. S2G**, and **Supplemental Dataset**).

We next conducted modified LEAP assays using the T4+AAddC oligonucleotide (**Table S1**). WT and EN-mutant, but not the RT-mutant, rORF2p could prime the synthesis of L1 cDNAs from RNA templates that ended in a 3’ poly(A) tail (**Fig. 1F**, bottom panel, JS+polyA and **Fig. S2E**, bottom panel, 3U+23A and 3U+polyA; and **Fig. S2F**, bottom panel, JS+23A) and to a much lesser extent from RNA templates lacking a poly(A) tail (**Fig. S2E**, bottom panel and **Fig. S2F**, bottom panel). Product characterization revealed that ddC was removed from the T4+AAddC oligonucleotide prior to its use as a primer in the LEAP reaction (**Fig. 1G**, **Fig. S2G**, and **Supplemental Dataset**).

To further investigate this unanticipated RT behavior, we tested a commercially available Moloney Murine Leukemia Virus reverse transcriptase (MMLV RT) enzyme in the LEAP reaction for comparison. The MMLV RT enzyme could use the T4+VN-3’-OH, but not the T4+AAddC, oligonucleotide to prime cDNA synthesis from RNA templates containing a 3’ poly(A) tail (**Fig. 1F** and **Fig. S2E**); no products were observed with RNA templates lacking a poly(A) tail (**Fig. 1F** and **Fig. S2E,** bottom panel). Unlike L1 ORF2p RT, which can prime reverse transcription in the absence of primer/template terminal complementarity (41,54,57), the MMLV RT cDNA products were all derived from a complementary primer/RNA template complex (**Fig. S2H** and **Supplemental Dataset**). Together, these data indicate that WT and EN-deficient rORF2p preparations contain a nuclease activity that can process a 3’ end modification from an oligonucleotide, which allows it to prime L1 cDNA synthesis. In contrast, MMLV RT does not possess such activity.

### WT and EN-rORF2p have an endonuclease activity that can remove an end modification from an exonuclease resistant oligonucleotide

To gain insight into the nature of the nuclease activity associated with L1 RNPs, immunoprecipitated ORF2p samples, and rORF2p samples, we designed an oligonucleotide that ends with three thymidine nucleosides, which are connected by phosphorothioate bonds, followed by an inverted thymidine residue (**Table S1**, T4+3T*invT, where the asterisk indicates the phosphorothioate bonds between the last three thymidine residues and the 3’ carbon of the penultimate deoxyribose sugar is connected to the 3’ deoxyribose sugar of the inverted thymidine [*i.e.*, forming a 3’-3’ linkage]). We reasoned that the T4+3T*invT oligonucleotide would both be resistant to 3’-5’ exonucleolytic degradation and block DNA synthesis, allowing us to gain insight into the type of nuclease activity present in our ORF2p preparations.

We first tested whether the T4+3T*invT oligonucleotide could be used to prime L1 cDNA synthesis using aliquots of crude RNPs (**Fig. 2A**) or immunoprecipitated ORF2p (**Fig. 2B**) in the LEAP reaction. WT (JJ101 and pTMO2F3) and EN-mutant (D145A and D205A), but not RT-mutant (D702A), samples efficiently primed the synthesis of L1 cDNAs using the T4+3T*invT primer (**Fig. 2A** and **Fig. 2B**). Product characterization from the immunoprecipitated ORF2p reactions revealed the 3’ end of the T4+3T*invT oligonucleotide likely was processed before being used to prime cDNA synthesis (**Supplemental Dataset** and see below).

**Fig. 2:**
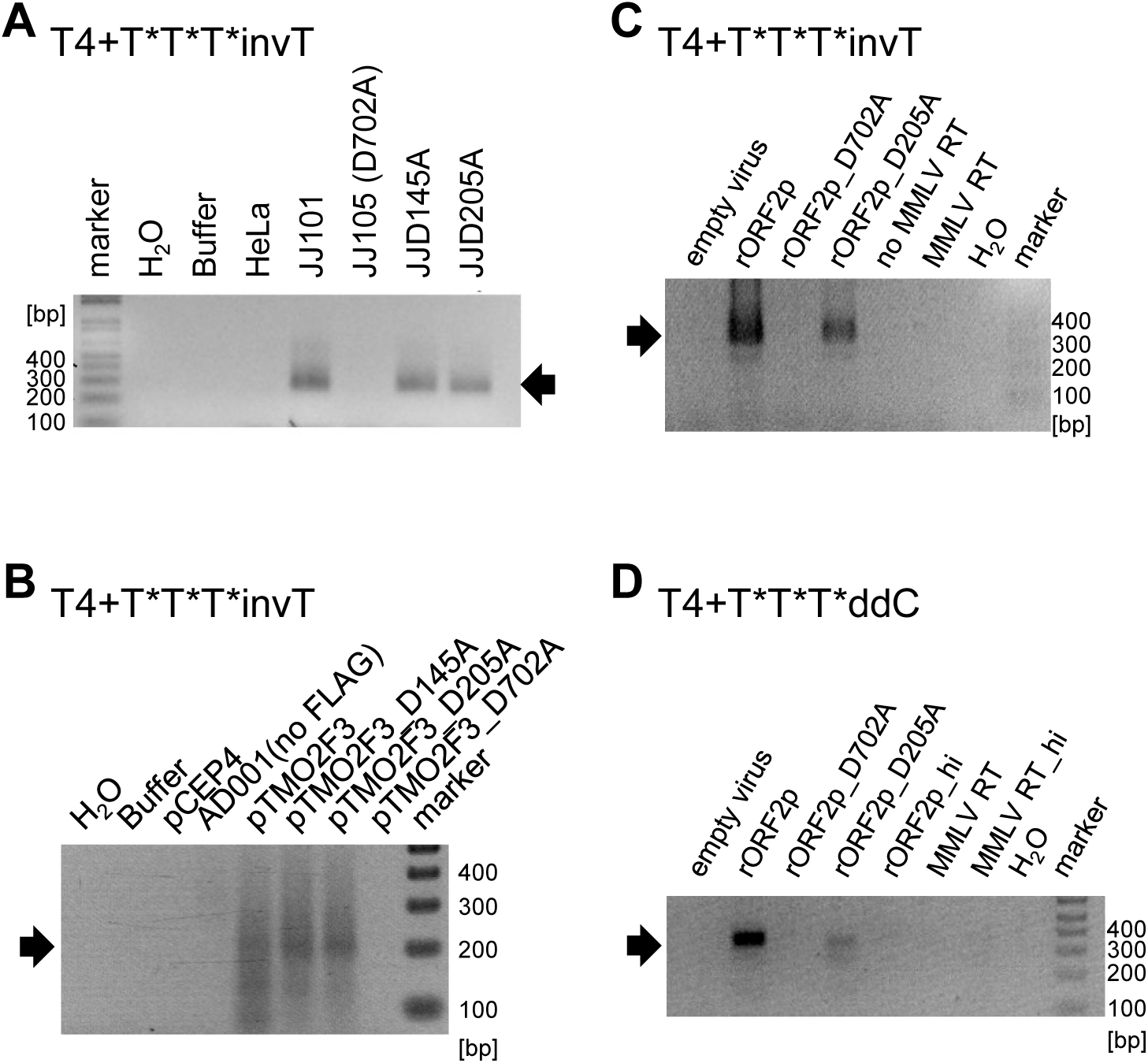
L1 RNPs, IP-ORF2p, and rORF2p can prime cDNA synthesis from exonuclease resistant oligonucleotides in a LEAP or modified LEAP reaction. LEAP (panels A and B) or modified LEAP reactions (panels C and D) were performed using exonuclease resistant oligonucleotides. The 3’-end of the oligonucleotides is indicated above each gel image. The asterisks indicate the position of phosphorothioate bonds in the oligonucleotide. The sample used in each reaction is indicated at the top of the gel image. Black arrows indicate the predicted size of the full-length products. Marker, molecular weight markers (bp). (*A*) *Results from LEAP reactions conducted using L1 RNPs and the T4+T*T*T*invT oligonucleotide*. LEAP reactions were conducted using RNPs derived from HeLa-JVM cells transfected with full length WT (JJ101), EN-mutant (JJD145A or JJD205A), or RT-mutant (JJ105/D702A). Negative controls: water (H_2_O, PCR control), buffer (LEAP control) and HeLa, (untransfected HeLa-JVM cells, extract control). Marker, molecular weight size controls (bp). At least three independent biological replicates were conducted for each reaction condition. *(B) Results from the LEAP assay using IP-ORF2p*. Immunoprecipitation reactions were conducted with whole cell extracts from HEK293T cells that were transfected with *L1.3 ORF2* expression constructs containing (pTMO2F3 constructs) or lacking (pAD001 no FLAG) an in-frame FLAG epitope tag at its 3’-end The controls and labeling are as described in Fig. 1B. pCEP4 is an empty vector control. pAD001 (no FLAG) serves as a negative control for the immunoprecipitation reaction. At least three independent biological replicates were conducted for each reaction condition. *(C and D) Results from the LEAP assay using rORF2p*. Modified LEAP reactions were conducted as described in Fig. 1F using either a T4+T*T*T*invT (panel C) or T4+T*T*T*ddC oligonucleotide (panel D). Negative controls: empty virus (no protein control), no MMLV RT (no protein control for MMLV RT reaction), and H_2_O (PCR control). MMLV RT (retrovirus control). In panel D, ‘hi’ indicates heat treated samples (rORF2p_hi and MMLV RT_hi) that serve as negative controls. Marker, molecular weight size controls (bp). At least three independent biological replicates were conducted for each reaction condition.

We next tested whether the T4+3T*invT oligonucleotide could be used to prime L1 cDNA synthesis in rORF2p samples using the modified LEAP assay (see **Fig. S2D**). The WT and EN-mutant, but not the RT-mutant, rORF2p samples efficiently promoted the synthesis of L1 cDNAs from an RNA template that contained a 3’ poly(A) tail (**Fig. 2C**, JS+polyA RNA; **Fig. S3A**, 3U+23A and 3U+polyA; and **Fig. S3B**, JS+23A), but not from an RNA template lacking a poly(A) tail (**Fig. S3A** and **Fig. S3B**). Similar results also were obtained using an oligonucleotide ending in T4+3T*ddC (**Fig. 2D** and **Table S1**). Product characterization revealed that the 3’ end of the T4+3T*ddC oligonucleotide was processed before being used to prime cDNA synthesis (**Supplemental Dataset**).

Additional experiments revealed that recombinant avian myeloblastosis virus reverse transcriptase (AMV RT) and MMLV RT could prime cDNA synthesis from an RNA template that contained a 3’ poly(A) tail using an oligonucleotide primer that ended in 12 thymidine residues and a 3’-OH group (**Table S1**, T12-3’-OH), but not from the T4+3T*invT oligonucleotide (**Fig. S3C**). Also, as opposed to the T4+VN-3’-OH and T4+AAddC oligonucleotides, the T4+3T*invT oligonucleotide was resistant to exonuclease I (ExoI) digestion (**Fig. S3D**), making it unlikely that a 3’ to 5’ exonuclease activity is responsible for removing the modification of from the T4+3T*invT oligonucleotide. Thus, we conclude that an alternative endonuclease (alt-EN) activity within our L1 RNP, immunoprecipitated ORF2p, and, most importantly, purified rORF2p samples can process the end modification from the T4+3T*invT oligonucleotide primer by cleaving internal phosphodiester linkages within the DNA chain.

### An RT-independent ligation-based assay to monitor alt-EN activity

The LEAP and modified LEAP assays require the production of a cDNA to detect L1 ORF2p RT activity, which prevents us to discern whether missense mutations within the L1 RT active site specifically affect alt-EN activity. To overcome this limitation, we designed a reverse transcription-independent assay to monitor the removal of a 3’ end modification from an oligonucleotide using a ligation-based readout that detects the cleavage of a 3’ group that is predicted to block the ligation of a 3’ end modified oligonucleotide (**Table S1**: *e.g.* T4+AAddC or T4+3T*invT) by assessing its ability to link to the 5’ phosphate of a second DNA molecule (**Fig. S4B**).

Briefly, we incubated aliquots of WT, EN-mutant, and RT-mutant rORF2p with an oligonucleotide that contains a 3’ end modification predicted to block DNA ligation (**Fig. 3A**, T4+AAddC or T4+3T*invT, indicated by the red slashed circle) and a 5’ biotin moiety (**Fig. 3A**, black circle labeled “B”) in a modified LEAP reaction condition lacking dNTPs (see Methods). After incubation, the oligonucleotide was captured using streptavidin beads and washed to remove rORF2p and any potentially contaminating proteins or nucleic acids (see **Methods**). We reasoned that alt-EN activity present within the rORF2p samples would process the blocking modification from the oligonucleotide, allowing the ligation of the resultant 3’-OH group to a second oligonucleotide containing a 5’-phosphate group (**Fig. 3A**, 5’-P blue line). A complementary “splint” or “bridging” oligonucleotide was used to facilitate the ligation reaction (**Fig. 3A**, purple line). Quantitative Taqman PCR reactions then were used to verify and quantify the presence of the ligated products (**Fig. 3A**, PCR primers, red arrows; Taqman probe, red table-top). The resultant data revealed a significant accumulation of ligation products relative to no protein and heat inactivated rORF2p controls when the T4+AAddC and T4+3T*invT oligonucleotides were incubated with WT, EN-mutant, but not the RT-mutant, rORF2p (**Fig. 3B**).

**Fig. 3:**
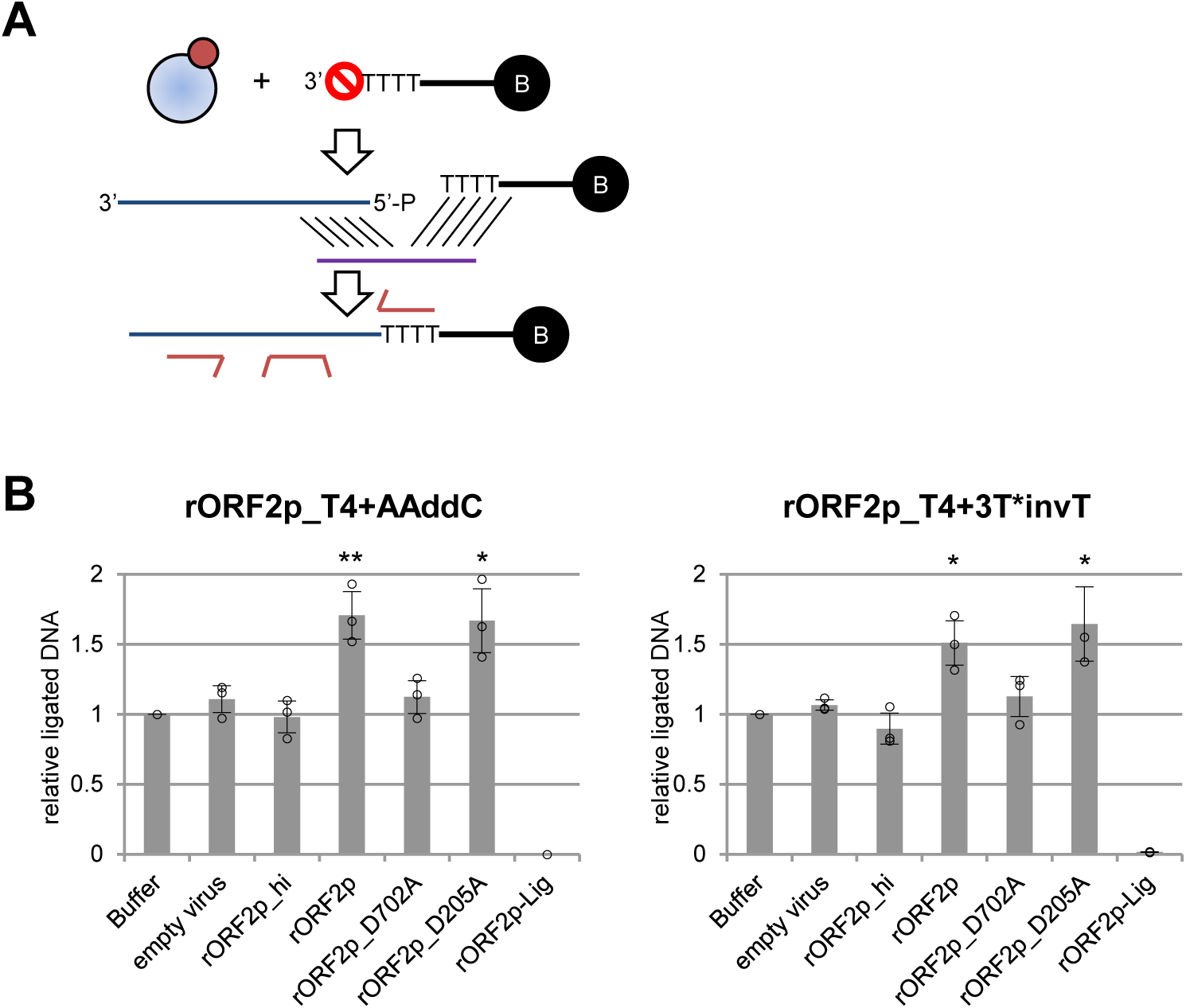
WT and EN-mutant, but not RT-mutant, rORF2p have alt-EN activity in a RT-independent ligation-based assay. *(**A**) Rationale of the assay.* WT and EN-deficient ORF2 proteins, but not a RT-deficient ORF2 protein, can remove a modification that blocks ligation from single strand DNA oligonucleotides. An aliquot of a rORF2p protein (blue/maroon circles) was incubated with an oligonucleotide containing a 3’ end modification predicted to block DNA ligation (red slashed circle) and a 5’ biotin moiety (black circle labeled with a B) in LEAP reaction buffer lacking dNTPs. After the reaction, the oligonucleotide was captured using streptavidin beads, washed, and tested for its ability to ligate to a second 5’ phosphorylated oligonucleotide (blue line) in the presence of a splint oligonucleotide (purple line). After streptavidin purification following the ligation reaction, the resultant products then were PCR amplified (red arrows). The tabletop symbol indicates the approximate position of the quenching oligonucleotide used in the Taqman PCR reactions. (***B*** *and **C***) *Results from the RT-independent ligation-based assay.* Reactions were conducted using oligonucleotides the ended in either T4+AAddC (panel B) or T4+3T*invT (panel C). The X-axis indicates the rORF2p sample used in the reaction. The Y-axis indicates the average amount of ligated DNA products detected when normalized to the buffer control. In panels B and C, ‘hi’ indicates samples treated with heat (rORF2p_hi) prior to conducting the RT-independent ligation-based assay. The rORF2p-Lig sample indicates a no ligase negative control. The error bars represent the standard deviation from three independent experiments. Statistical significance was gauged relative to the buffer control using the Student’s t-test (*P < 0.05; **P < 0.01).

Additional controls, using an oligonucleotide ending in 4 thymidine residues and a 3’-OH group (T4-3’-OH), showed that the detection of ligation products required the T4-3’-OH oligonucleotide, a 5’-phosphorylated acceptor oligonucleotide, and the “splint” oligonucleotide (**Fig. S4A**). Moreover, the addition of RNA to the reactions did not enhance ligation efficiency using the T4+AAddC and T4+3T*invT oligonucleotides (**Fig. S4C**). Notably, we did not observe differences in alt-EN activity among the WT, EN-mutant, and RT-mutants when the RT-independent ligation-based assays were conducted with crude RNP samples (**Fig. S5A**, **Fig. S5B**, and **Fig. S5C**). In contrast, the differences in alt-EN activity, albeit modest, were observed when immunoprecipitated ORF2p samples were used in this RT-independent ligation-based assay (**Fig. S5D**, **Fig. S5E**, and **Fig. S5F**), suggesting that rORF2p harbors alt-EN activity; however, we cannot rule out the possibility that similar cellular activities may be present in the L1 RNP and IP-ORF2p samples.

### Different rORF2p preparations produced in insect cells have alt-EN activity

The results from the RT-independent ligation-based assays suggest that WT and EN-mutant, but not RT-mutant, rORF2p contain alt-EN activity. However, we could not rule out the possibility that a low-level contaminating nuclease activity within the rORF2p preparations is responsible for the observed alt-EN activity. To address this concern, we purified rORF2p using an additional polyethyleneimine (PEI) precipitation step (62) to remove nucleic acids and nucleic acid binding proteins from the rORF2p preparations; we also included extensive wash steps to remove potential contaminants and a non-ionic non-denaturing detergent (IGEPAL CA-630) from rORF2p preparations (**Fig. S6A**, **Fig. S6B**, and **Fig. S6C**, see Methods).

Aliquots from the three WT rORF2p preparations (rORF2p+IGEPAL-PEI [used above], rORF2p+IGEPAL+PEI, and rORF2p-IGEPAL-PEI) then were tested for their ability to use different oligonucleotides (**Table S1**: T4+VN-3’-OH, T4+AAddC, T4+3T*invT, and T4+3T*ddC) to prime cDNA synthesis from a JS+23A RNA template using a modified LEAP reaction. Each WT rORF2p sample could use the T4+VN-3’-OH, T4+AAddC, T4+3T*invT, and T4+3T*ddC oligonucleotides to prime the reverse transcription of the JS+23A RNA template (**Fig. 4A**). The EN-mutant, but not RT-mutant, rORF2p, which were prepared in either the presence (+) or absence (-) of a PEI precipitation step, could use the T4+VN-3’-OH, T4+AAddC, T4+3T*invT oligonucleotides to prime cDNA synthesis from a JS+23A RNA template (**Fig. S6D**). Product characterization revealed that the 3’ end modification was removed from the T4+AAddC and, likely, T4+3T*invT oligonucleotides prior to their use as primers in cDNA synthesis (**Supplemental Dataset**).

**Fig. 4:**
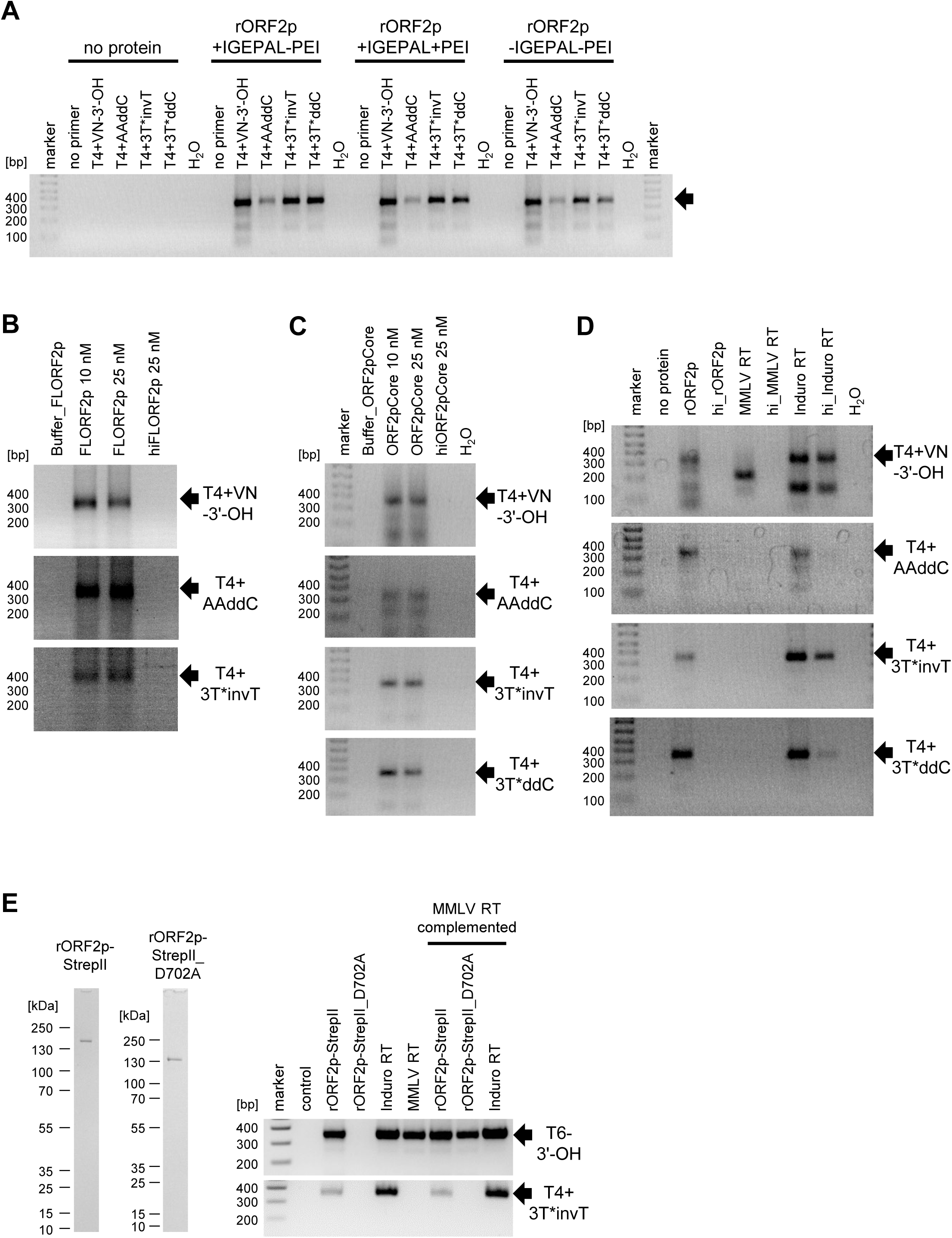
ORF2 proteins purified using different protocols and a thermostable group II intron-encoded RT have alt-EN activity. *(**A**) Results of the modified LEAP assay.* Recombinant ORF2p was purified with or without an additional polyethyleneimine precipitation step (+PEI or -PEI, respectively). In one sample, an additional wash was included to remove the detergent (IGEPAL CA-630) from the sample (see **Materials and Methods**). Aliquots of the resultant samples (rORF2p+IGEPAL-PEI, rORF2p+IGEPAL+PEI, and rORF2p-IGEPAL-PEI) were subjected to a modified LEAP reaction using the T4+VN-3’-OH, T4+AAddC, T4+3T*invT, or T4+3T*ddC to prime cDNA synthesis from a JS+23A RNA template. Oligonucleotide names are indicated at the top of the gel image. Negative controls: H_2_O (PCR control), no primer (modified LEAP control). Marker, molecular weight standards in bp. The expected size of the modified LEAP product is indicated by the black arrow. At least two independent biological replicates were conducted for each purified sample. *(**B** and **C**) Recombinant ORF2p purified from insect cells using a different protocol and a bacterially produced ORF2p have alt-EN activity*. Full-length ORF2p (panel B) and an rORF2p “core” protein lacking the L1 ORF2 EN and C domains (panel C) were generous gifts from Dr. Martin S. Taylor and Dr. Trevor van Eeuwen. Aliquots of each protein were used in modified LEAP reactions to test whether the indicated oligonucleotides (right of the gel images) could prime cDNA synthesis from a JS+23A RNA template. The molar concentration of each protein is indicated at the top of the gel image. Negative controls: buffer (no protein control), hi ORF2p (heat inactivated protein), and H_2_O (PCR control). Marker, molecular weight standards in bp. The expected size of the modified LEAP product is indicated by the black arrow. Each experiment with one biologically independent sample was technically repeated at least three times. *(**D**) A commercially available thermostable group II intron-encoded protein has alt-EN activity.* Induro RT was used in a modified LEAP assay to test whether the indicated oligonucleotides (right of gel image) could prime cDNA synthesis from a JS+23A RNA template. Positive control (rORF2p). Negative controls: no protein, heat inactivated rORF2p (hi_rORF2p) or MMLV (hi_MMLV RT). Note: heat inactivation reduced but did not abolish Induro activity; MMLV could only use the T4+VN-3’-OH oligonucleotide to prime cDNA synthesis. The expected size of the modified LEAP product is indicated by the black arrow. At least three independent replicates were conducted for each reaction condition. *(**E**) A missense mutation in the L1 rORF2p RT active site eliminates alt-EN activity*. A purified version of rORF2p containing an in-frame Strep-II epitope tag at its carboxyl terminus (rORF2p-StrepII) (Left panel) was used in modified LEAP assays (Right panel) with either a T6-3’-OH (top) or T4+3T*invT primer (bottom) to prime cDNA synthesis from the JS+polyA RNA template. MMLV RT complemented indicates that MMLV RT was mixed with the indicated proteins (three right gel lanes). Note: the rORF2p-StrepII_D702A mutant + MMLV RT reaction lacked discernable products using the T4+3T*invT primer when compared to the rORF2p-StrepII + MMLV RT and Induro RT + MMLV RT control reactions (bottom gel panel, far right lanes), suggesting that the L1 ORF2p RT D702A mutation eliminates both L1 cDNA synthesis and alt-EN activity. Negative control: buffer only. Marker, molecular weight standards in bp. The expected size of the modified LEAP product is indicated by the black arrow. Three independent replicates were conducted for each reaction condition.

We next used an aliquot of full-length L1 ORF2p that was purified from insect cells using FLAG affinity chromatography, heparin affinity chromatography, and size exclusion chromatography (a gift from Dr. Martin S. Taylor and Dr. Trevor van Eeuwen), which recently was used to determine a cryogenic electron microscopy (Cryo-EM) structure of L1 ORF2p (26). As above, this protein, FLORF2p, could use the T4+VN-3’-OH, T4+AAddC, T4+3T*invT oligonucleotides to prime cDNA synthesis from a JS+23A RNA template and product characterization confirmed that the 3’ end modification was removed from the T4+AAddC, T4+3T*invT oligonucleotides prior to their use as primers in the modified LEAP reaction (**Fig. 4B**, left panel and **Supplemental Dataset)**. Thus, we conclude that WT and EN-mutant rORF2p prepared using different purification schemes contain alt-EN activity.

### An rORF2p “core” protein produced in E. coli has alt-EN activity

The above data strongly suggest the WT and EN-mutant rORF2p have alt-EN activity. However, because each of these proteins was purified from insect cells, it remained possible that an occult contaminant from insect cells could be responsible for alt-EN activity. To examine this possibility, we obtained a rORF2p “core” protein that was extensively purified from *E. coli* using nickel-nitrilotriacetic acid (Ni-NTA) affinity chromatography, heparin affinity chromatography, and size exclusion chromatography (a generous gift from Dr. Martin S. Taylor and Dr. Trevor Van Eeuwen) and recently was used to determine the X-ray crystallographic structure of the L1 ORF2p RT domain (26). Notably, the ORF2p “core” lacks the EN and cysteine-rich (C) domains.

Modified LEAP reactions revealed that the rORF2p “core” could use the T4+VN-3’-OH, T4+AAddC, T4+3T*invT, and T4+3T*ddC oligonucleotides to prime cDNA synthesis from a JS+23A RNA template (**Fig. 4C**). Product characterization confirmed that the 3’ end modification was removed from the T4+AAddC, T4+3T*invT, and T4+3T*ddC oligonucleotides before their use as primers to initiate cDNA synthesis (**Supplemental Dataset**). Thus, rORF2p proteins isolated from insect and bacterial cells have alt-EN activity.

### A commercially available group II intron-encoded protein has alt-EN activity

Bacterial mobile group II introns encode ancient reverse transcriptase enzymes that are proposed to be the evolutionary progenitor of the RT domains present in non-LTR retrotransposons, telomerase, and modern retroviruses (reviewed in (63,64) also see (65)). Moreover, some mobile group II intron-encoded RTs lack an endonuclease domain and can exploit a 3’-OH group present on Okazaki fragments to mediate their mobility by an EN-independent retrotransposition-like pathway (66). Given the close phylogenetic relationship between the group II intron and L1 ORF2p RT domains, we next sought to examine whether a commercially available thermostable group II intron-encoded RT that lacks an endonuclease domain (Induro RT) contains alt-EN activity.

An aliquot of Induro RT (New England BioLabs) was used in a modified LEAP assay to examine whether it allowed the T4+VN-3’-OH, T4+AAddC, T4+3T*invT, and T4+3T*ddC oligonucleotides to prime cDNA synthesis from a JS+23A RNA template. We readily detected Induro RT alt-EN activity using each oligonucleotide at both 37°C (**Fig. 4D** and **Fig. S6E**) and 55°C (**Fig. S6F**, the optimal reaction temperature for Induro RT). Product characterization revealed that the lower molecular weight product in Induro RT reactions using the T4+VN-3’-OH primer (**Fig. 4D** and **Supplemental Dataset**) is due to the use of upstream priming sites within JS+23 RNA template, that Induro RT could initiate reverse transcription in the absence of primer/template terminal complementarity, and that 3’ end modifications were removed from the T4+AAddC, T4+3T*invT, and T4+3T*ddC oligonucleotides prior to their use as primers for cDNA synthesis (**Fig. S6G**, **Fig. S6H,** and **Supplemental Dataset**). Notably, pretreating Induro RT at 90°C for 20 minutes did not completely inactivate alt-EN or RT activity (**Fig. 4D** and **Fig. S6E**) — a treatment that should eliminate any contaminating nucleases in the Induro preparation. By comparison, pretreating Induro RT with protease K abolished both alt-EN and RT activity (**Fig. S6E**). In sum, these data provide strong evidence that Induro RT has both RT and alt-EN activities.

### A mutation in the L1 ORF2p RT active site eliminates alt-EN activity

To further confirm that the L1 ORF2p RT domain is responsible for alt-EN activity, we further purified a version of rORF2p containing a Strep-II epitope tag at its carboxyl terminus (rORF2p-StrepII) and conducted modified LEAP assays using either a T6-3’-OH or T4+3T*invT primer (**Fig. 4E**). The rORF2p-StrepII, Induro RT and MMLV RT, but not the RT-deficient rORF2p-StrepII_D702A mutant, were able to synthesize cDNAs using the T6-3’-OH primer (**Fig. 4E**, top panel). In contrast, only rORF2p-StrepII and Induro RT could promote cDNA synthesis using the T4+3T*invT primer (**Fig. 4E**, bottom panel).

We next mixed the rORF2p-StrepII, rORF2p-StrepII_D702A RT-mutant, and Induro RT proteins with MMLV RT. We reasoned that if a contaminating nuclease in our preparations was responsible for processing the 3’-end of the T4+3T*invT primer, the MMLV RT protein would be able to use the processed T4+3T*invT primer to synthesize cDNAs, resulting in positive LEAP signals in all reactions regardless of whether WT or RT-mutant rORF2p was used in the experiment. Notably, the rORF2p-StrepII + MMLV RT and Induro RT + MMLV RT reactions supported cDNA synthesis, whereas the rORF2p-StrepII_D702A RT-mutant + MMLV RT reaction lacked discernable products using the T4+3T*invT primer (**Fig. 4E**, bottom panel). Thus, these data strongly suggest that L1 ORF2p and Induro RT proteins can remove a 3’ block from a DNA primer, enabling subsequent primer extension. A missense mutation in the L1 ORF2p RT active site eliminates alt-EN activity, implying that this activity is contained within the ORF2p RT.

## Discussion

We previously observed that RNPs derived from cells transfected with WT and EN-deficient mutant L1 constructs contain an nuclease activity that could process a 3’ modification in a single strand oligonucleotide primer that blocks DNA synthesis, allowing it to be used to prime L1 cDNA synthesis; however, we could not definitively determine whether this activity was contained within L1 ORF2p or was due to a co-isolated cellular nuclease in our RNP preparations (54). Here, we used several different forms of purified ORF2p and developed both modified LEAP and RT-independent ligation-based assay to determine whether this nuclease activity is contained within L1 ORF2p. Our data demonstrate: (**i**) recombinant full-length versions of ORF2p produced in insect cells using various protocols have alt-EN activity (**Fig. 1B**, **Fig. 1F**, **Fig. 2**, **Fig. S2D**, **Fig. S2E**, **Fig. S2F**, **Fig. 3**, **Fig. S3**, **Fig. 4A**, **Fig. 4B**, **Fig. 4E**, **Fig. S6A**, **Fig. S6B**, **Fig. S6C**, and **Fig. S6D**); (**ii**) a bacterially-expressed rORF2p core protein that lacks the ORF2p EN and C domains contains alt-EN activity (**Fig. 4C**) (26); and (**iii**) a commercially available thermostable group II intron-encoded protein (Induro RT), which, as best as we can determine, was expressed in bacteria and lacks an EN domain, contains alt-EN activity (**Fig. 4D**, **Fig. 4E**, **Fig. S6E**, and **Fig. S6F)**; and (**iv**) the ORF2p RT active site is required for alt-EN activity and a D702A mutant eliminates primer 3’ end processing (**Fig. 3B**, **Fig. 4E**, **Fig. S5E**, and **Fig. S5F**). Together, these data make a compelling case that L1 ORF2p, and the Induro group II intron-encoded protein, contain an evolutionarily conserved alt-EN activity within their RT modules.

Critical to our analyses was the use of different oligonucleotide primers to examine RT and alt-EN activity in modified LEAP assays. Importantly, IP-ORF2p, full-length rORF2p, the bacterially-expressed rORF2p core protein, and the group II intron-encoded Induro RT protein could process a 3’-5’ exonuclease I resistant blocking lesion from either the T4+3T*invT or T4+3T*ddC oligonucleotides (**Fig. S3D**) before they could be used to prime reverse transcription in our modified LEAP assays (**Fig. 2**, **Fig. S3A**, **Fig. S3B**, **Fig. 4**, **Fig. S6D**, **Fig. S6E**, and **Fig. S6F**). By comparison, commercially available retroviral RTs (MMLV and AMV) contained severely reduced alt-EN activity under our reaction conditions (**Fig. 1F**, **Fig. 2C**, Fig**. 2D**, **Fig. S2E**, **Fig. S3A**, **Fig. S3C**, **Fig. 4D**, **Fig. 4E**, and **Fig. S6F**). Notably, IP-ORF2p and full-length rORF2p also could process the T4+3T*invT primer used in our RT-independent ligation-based assay (**Fig. 3**, **Fig. S4**, and **Fig. S5**) and complementation assay (**Fig. 4**).

Characterization of the products from our modified LEAP reactions yielded insights about the RT and alt-EN activities contained within the various RT proteins examined in our study. We found that full-length ORF2p (IP-ORF2p and rORF2p), the bacterially expressed rORF2p core protein, and Induro RT could initiate reverse transcription from oligonucleotide template/RNA substrate complexes that lack perfect terminal complementarity (**Fig. 1C**, **Fig. 1G, Fig. S2G**, **Fig. S2H**, **Fig. S6G**, **Fig. S6H**, and **Supplemental Dataset**). In contrast, MMLV RT could only initiate reverse transcription from oligonucleotide template/RNA substrate complexes containing perfect complementarity at their 3’ ends (**Fig. 1G**, **Fig. S2H**, and **Supplemental Dataset**).

Our RT-independent ligation-based assay revealed that WT ORF2p and EN-deficient mutant versions of ORF2p, but not an ORF2p RT-deficient (D702A) mutant, can remove 3’ blocking groups from the T4+AAddC and T4+3T*invT oligonucleotides, thereby allowing them to be ligated to a 5’ phosphorylated oligonucleotide (**Fig. 3**). The findings that a bacterially expressed rORF2p core protein that lacks the ORF2p endonuclease and cysteine rich domains contains alt-EN activity (**Fig. 4C**), and that a rORF2p D702A RT-deficient mutant severely inhibits the ability to remove a 3’ blocking group from the T4+3T*invT oligonucleotide in complementation assays conducted in the presence of MMLV RT (**Fig. 4E**), imply that the ORF2p RT active site is required for both L1 cDNA synthesis (a polymerization/nucleophilic addition reaction) and alt-EN activity (a hydrolysis reaction).

Previous studies have made a strong case that the RT domains of group II intron-encoded proteins, LINE elements, telomerase, and retroviruses are derived from a primordial reverse transcriptase (**Fig. 5A**, RT origin) that likely played a critical role in the transition from an “RNA” to a “DNA” world (*e.g.*, see (67–70)). Of note, telomerase also is associated with a “proofreading” nuclease activity that can remove non-telomeric nucleotides from single strand telomeric DNA repeats before initiating the telomere extension (*i.e.*, cDNA synthesis) from a complementary telomeric DNA primer/telomerase RNA template complex (71–73), which is similar to the alt-EN activity reported in our study, further highlighting similarities between the L1 ORF2p RT activity and telomerase (54).

**Figure 5.**
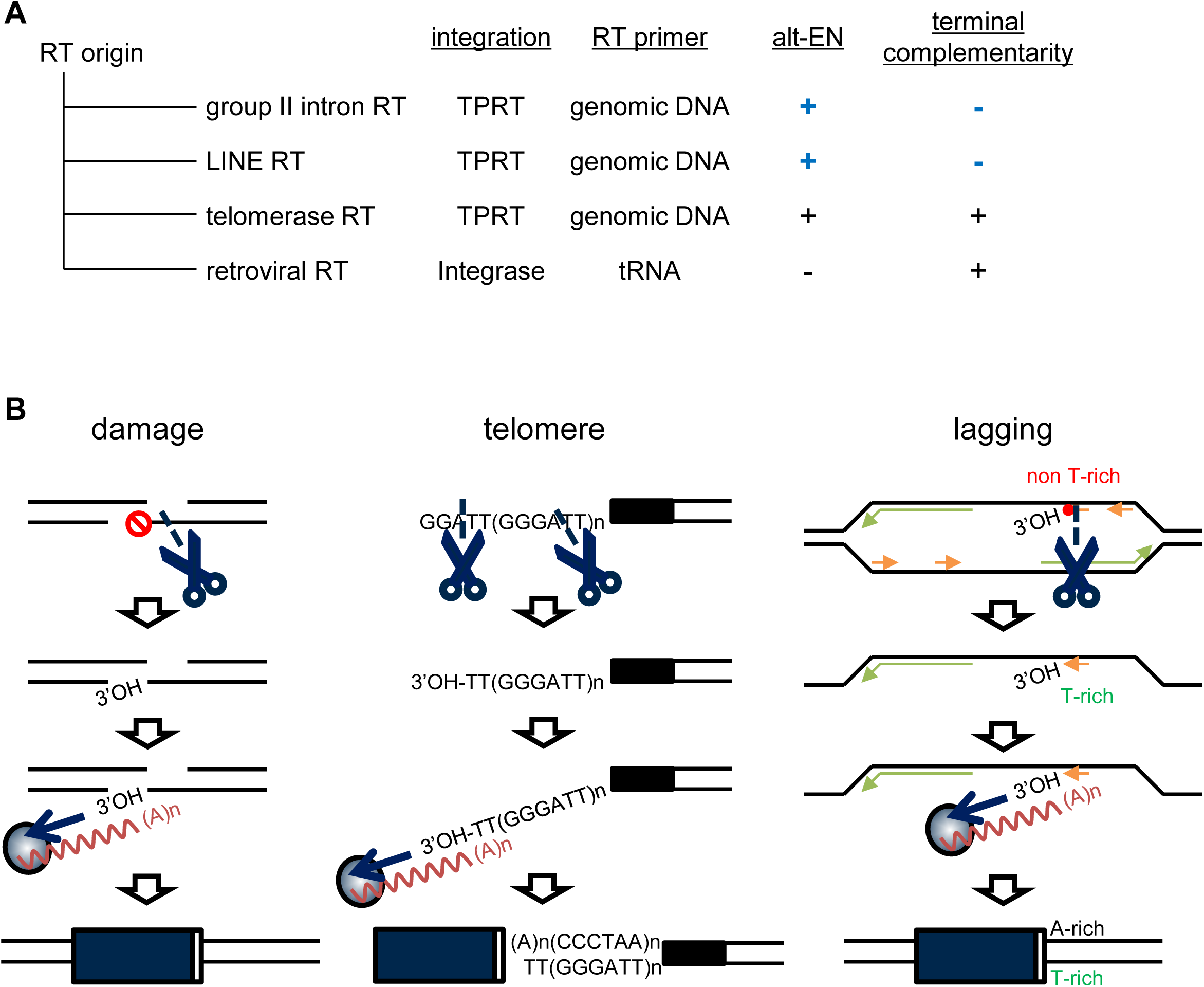
Evolution and proposed biological roles of alt-EN nuclease activity. ***(A)*** *Relatedness of the RT proteins encoded by different retroelements*. Column 1, a primordial reverse transcriptase (RT origin) likely gave rise to the RTs encoded by group II introns, LINEs, telomerase, and retroviral RTs. Column 2, the mode of integration used by each retroelement in Column 1. Column 3, the source of the primer used to initiate reverse transcription. Column 4, whether the protein contains alt-EN activity. Column 5, whether the initiation of reverse transcription requires terminal complementarity between the DNA primer and RNA template. Blue lettering, results from this study. Black lettering, results of previous studies. *(**B**) Possible roles for alt-EN in endonuclease-independent L1 retrotransposition*. Left, The L1 ORF2p alt-EN activity may remove a modification that blocks cDNA synthesis from the site of an endogenous genomic DNA lesion (red circle with line), exposing a 3’-OH group. The resultant 3’-OH group then may allow the L1 ORF2p RT (blue circle) to initiate L1 cDNA synthesis (blue arrow) from the L1 RNA template (red wavy line). Center, regions of micro-complementarity between the L1 RNA 3’ poly(A) tail and single strand telomeric DNA repeat may allow the ORF2p (blue circle) alt-EN activity create an endonucleolytic nick in single strand telomeric DNA at L1 RNA/telomeric DNA hybrids. The cleavage then would expose a short stretch of DNA that ends in one or two thymidine residues and terminates in a 3’-OH group. The resultant 3’-OH group then may allow the L1 ORF2p RT to initiate cDNA synthesis (blue arrow) from the L1 RNA template (red wavy line). Right, L1 ORF2p alt-EN may remove a 3’ modification from an Okazaki fragment present at a stalled replication fork (red circle), thereby allowing it to serve as a substrate for ENi retrotransposition.

Considering our results in a broader perspective, and assuming a monophyletic origin of reverse transcriptases (70), we speculate that the primordial reverse transcriptase (**Fig. 5**, RT origin), and perhaps the RNA-dependent RNA polymerase from which it many have been derived, also may have contained alt-EN activity. We further propose that this activity was dampened or lost in retroviral reverse transcriptases, which initiate reverse transcription from a transfer RNA primer/retroviral RNA template complex with perfect complementarity at the primer 3’ end. It will be interesting to determine whether the loss of specific subdomains in retroviral RTs, which are present in retroelements that initiate cDNA synthesis using TPRT, play a role in the loss of alt-EN activity (**Fig. 5A**) (65).

The finding that L1 RT and Induro RT confer alt-EN activity, in addition to their documented function in RNA-templated DNA synthesis, highlights an unanticipated flexibility of these enzymes with ramifications for mobile genetic element biology. Although most enzymes are remarkably specific catalysts, some can execute different reactions using a single active site (74). Examples include lipases that catalyze the hydrolysis of diverse lipidic substrates, but also can promote the transfer of an acyl group from lipids to cholesterol (*i.e.*, a transferase activity), as well as Glutathione S-transferases, cytochrome P450, and serum paraoxonase, which can act on a broad range of dissimilar molecules (75,76). Intriguingly, RNA polymerases, which add ribonucleotides to an RNA chain, also can use the same active site to hydrolyze terminal nucleotides to correct misincorporation errors during transcription (77–80). In contrast, DNA polymerases, which like RTs catalyze the synthesis of deoxyribonucleotide chains, typically perform proofreading activities using separate nuclease domains or the exonucleolytic activity of auxiliary factors. (81,82). How the compact L1 RT active site can accommodate different nucleic acid substrates for both synthesis and hydrolysis require further exploration.

How might alt-EN function in L1 retrotransposition? It has been postulated that enzyme promiscuity serves as starting point for the evolution of new functions, and adaptive evolution experiments support this idea (83,84). For L1 RT, it is likely that alt-EN is a primordial function that was critical in ancestral retrotransposons prior to the acquisition of the *L1 ORF2p* EN and cysteine-rich domains (45). Thus, we propose that alt-EN activity may be critical to remove blocking groups from the 3’ ends of genomic DNA substrates, thereby allowing L1 to integrate at spontaneous DNA lesions (52,53), dysfunctional telomeres (55), or stalled DNA replication forks (45) via ENi retrotransposition (**Fig. 5B**). Indeed, DNA break sites often have terminal damage or synthesis incompatible ends (85,86), requiring specialized mechanisms to ‘clean’ DNA ends before cells can repair or re-synthesize broken DNA. We speculate that the flexibility of the L1 RT active site, coupled with an inherent mechanism to process these ends (*i.e.*, alt-EN activity) for use in retrotransposon integration, may have greatly expanded opportunities for retrotransposon dissemination. In sum, our study provides novel insights into the mechanisms used to initiate reverse transcription by TPRT-like reactions and sheds light on the evolution of retrotransposons and retroviruses.

## ACKNOWLEDGEMENTS

We thank Dr. Martin S. Taylor and Dr. Trevor van Eeuwen for providing highly purified versions of FLORF2p and L1 ORF2p “core” protein. We thank Lionel Jond for technical assistance with baculovirus production. We also thank members of the Moran and Barabas laboratories, as well as members of the University of Michigan Genomic Instability group, for helpful discussions during the course of this study. Finally, we thank Dr. John B. Moldovan and Ms. Sierra Mortimer for critically reading this manuscript.

## AUTHOR CONTRIBUTIONS

M.N., H.C.K, M.D., O.B and J.V.M. designed the study. M.N., H.C.K., and M.D. conducted experiments. M.N., H.C.K, M.D., O.B and J.V.M. analyzed data. M.N., H.C.K, M.D., O.B and J.V.M. wrote and edited the manuscript.

## DATA AVAILABILITY

All the data underlying the findings in this manuscript are included within this submission.

