## Supplemental data for "An Alternative DNA Endonuclease Activity is Associated with the LINE-1 ORF2-encoded Protein"

#### Supplemental Figure Legends and Table

**Figure S1: LEAP assays conducted with L1 RNPs and immunoprecipitated ORF2p**. ***(A)*** *Rationale of the assays*. L1 RNPs (left) or immunoprecipitated ORF2p (IP-ORF2p) were incubated with an oligonucleotide that ends in a 3' polythymidine tract (TTTT-3'OH; black line) and dNTPs. Base pairing between the 3' end of the oligonucleotide and the 3' poly(A) tract of L1 RNA (wavy black line) allows the formation of an oligonucleotide primer/L1 RNA template complex. The ORF2p RT activity can use the resultant primer/template complex to synthesize L1 (-) strand cDNA (blue arrow). The resultant cDNA can be amplified using primers specific for the oligonucleotide primer (purple arrow) and L1 RNA (red arrow). ***(B)*** *Flow chart detailing the preparation of L1 RNPs.* HEK293T cells transfected with an engineered WT L1 (JJ101/L1.3), RT-deficient (JJ105/L1.3 [D702A]), or EN-deficient mutant (JJD145A or JJD205A) L1 constructs were subjected to Hygromycin B selection. Whole cell lysates (WCLs) prepared from the hygromycin-resistant cells and then were subjected to ultracentrifugation through a sucrose cushion to isolate L1 RNPs (41,61) (see **Materials and Methods**). *(**C**) Western blot analyses reveal ORF1p is enriched in L1 RNPs*. Left, western blots using 30 µg of WCLs. Right, western blots using 3.5 µg of L1 RNPs. The engineered L1 constructs transfected into HEK293T cell are indicated at the top of the gels. Antibodies used to detect ORF1p (hORF1N), ribosomal protein S6 (S6), and β-actin (loading control) are indicated at the left of the gel image. Molecular weight markers (kDa) are shown at the right of the gel image. pCEP4 is an empty vector control and served as a negative control. AJ101 is a monocistronic L1 construct that expresses ORF2p but lacks *L1 ORF1* coding sequences and served as a negative control. At least three independent biological replicates were conducted for each purified sample. *(**D**) Flow chart detailing the preparation of IP-ORF2p*. HEK293T cells were transfected with a monocistronic ORF2p expression construct containing three copies of an in-frame FLAG epitope tag (pTMO2F3), pTMO2F3 EN-deficient mutant constructs (pTMO2F3_D145A or pTMO2F3_D205A) or a pTMO2F3 RT-deficient mutant construct (pTMO2F3_D702A). The transfected cells were subjected to Hygromycin B selection. Whole cell lysates (WCLs) prepared from the hygromycin-resistant cells then were subjected to immunoprecipitation using an anti-FLAG antibody to isolate IP-ORF2p (58) (see **Materials and Methods**). *(**E**) Western blot analyses reveal ORF2p is enriched in IP-ORF2p* *preparations*. Left, western blots using 30 µg of WCLs. Right, western blots using 3.5 µl of the resultant IP-ORF2p solution. The engineered L1 constructs transfected into HEK293T cell are indicated at the top of the gels. Antibodies used to detect L1 ORF2p (FLAG), ribosomal protein S6 (S6), and β-actin (loading control) are indicated at the left of the gel image. Molecular weight markers (kDa) are shown at the right of the gel image. pCEP4 is an empty vector control and served as a negative control. AD001 is a monocistronic ORF2p expression construct that lacks the FLAG epitope tag and served as a negative control. At least three independent biological replicates were conducted for each purified sample.

**Figure S2: Purification of rORF2p**. *(**A**) The rORF2p purification protocol.* High Five cells were infected with a baculovirus encoding a recombinant human L1.3 ORF2p containing an in-frame carboxyl-terminal 3xFLAG/3xHA epitope tag. After three days of incubation, the cells were lysed and rORF2p was purified using: (**i**) anti-FLAG affinity chromatography, (**ii**) anti-HA affinity chromatography, and (**iii**) heparin column chromatography. The eluate then was dialyzed to obtain purified rORF2p (see **Materials and Methods**). *(**B**) Western blot analyses of rORF2p*. The purified rORF2p samples were detected by western blot with anti-FLAG antibody (left), an anti-HA antibody (middle), and anti-hORF2N antibody (right). At least three independent biological replicates were conducted for each purified sample. *(**C**) In vitro synthesized RNAs*. All the RNA templates were synthesized using a T7 Quick High Yield RNA Synthesis Kit or HiScribe T7 High Yield RNA Synthesis kit and then were gel purified (see **Materials and Methods**). The sequence of the 3U and JS RNA was derived from the L1 3'UTR sequence present in pJCC5/L1.3 or the SV40 poly(A) signal present in JJ101/L1.3, respectively. The +23A and +polyA indicates RNA templates containing a 23 nucleotide 3' poly(A) tract or a 3' poly(A) tail, respectively. The purified RNA was separated on a denaturing polyacrylamide gel and visualized by GelRed staining solution. At least three independent biological replicates were conducted for each purified sample. *(**D**) Rationale of the modified LEAP assay using rORF2p*. The purified rORF2p (blue circle with dark red circle) was incubated with an *in vitro* synthesized RNA template (black wavy line), a single strand DNA oligonucleotide primer (TTTT-3'OH; black line), and dNTPs. The recombinant proteins then were tested for their ability to synthesize cDNA (blue arrow) from the DNA oligonucleotide/*in vitro* synthesized exogenous RNA template complexes. The resultant cDNAs were amplified using primers specific for the oligonucleotide primer (purple arrow) and *in* vitro transcribed RNA template (red arrow). ***(E** and **F**) RT activity of the purified rORF2p using the T4+VN-*3'-OH *(MN440) (top gel images) and T4+AAddC (MN488) (bottom gel images) oligonucleotides (**Table S1**) with exogenous RNAs*. The 3U, 3U+23A, and 3U+polyA *in vitro* transcribed RNAs were used as templates in panel E. The JS and JS+23A *in vitro* transcribed RNAs were used as templates in panel F. The rORF2p used in the modified LEAP reaction is indicated above each lane of the gel images. Reactions conducted with MMLV RT served as an additional control. The empty virus (no rORF2p), $H_2O$, no MMLV RT, and heat inactivated rORF2p (rORF2p_hi) served as negative controls. The black arrows indicate the expected length of the full-length RT-PCR products. At least three independent biological replicates were conducted for each condition. ***(G**) Characterization of the modified LEAP products*. We sequenced the modified LEAP products generated in **Fig. 1F**, **Fig. S2E**, and **Fig. S2F**. The protein samples used in the modified LEAP reaction are indicated at the left of the Table. The number of products sequenced from LEAP reactions using the *in vitro* transcribed RNAs (column 2) and the T4+VN-3'-OH and T4+AAddC oligonucleotide primers and are indicated at the top of the Table. The number of LEAP products (denominator) exhibiting a 3'-end modification (numerator) from each primer/template complex and the inferred 3'-end used to prime L1 cDNA synthesis (indicated below the primer names). ***(H**) Characterization of modified LEAP products using the T4+VN-3'-OH primer either the JS+23A or JS+polyA in vitro transcribed RNA templates from **Fig. 1F** and **Fig. S2F***. We only analyzed the products that initiated cDNA synthesis within the 3' polyA tail. The protein samples used in the modified LEAP reaction are indicated at the top the Table. The sequence of the “V” and “N” nucleotides in the modified LEAP products are indicated in the Table. Note: consistent with our previous analyses (41,54) rORF2p can extend primer/template complexes that lack perfect complementarity, whereas MMLV RT cannot.

**Figure S3: Purified rORF2p can initiate reverse transcription using an exonuclease-resistant oligonucleotide**. ***(A** and **B**) WT (rORF2p) and EN-deficient (rORF2p_D205A) proteins can initiate reverse transcription from an oligonucleotide primer that is blocked at its 3' end.* The sequence of the T4+3T*invT oligonucleotide primer is indicated in **Table S1** (MN503). Asterisks indicate the presence of phosphorothioate bonds between the last three thymidine residues; the 3' carbon of the penultimate deoxyribose sugar is connected to the 3' deoxyribose sugar of the inverted thymidine (*i.e.*, forming a 3'-3' linkage). *In vitro* transcribed RNAs used as templates in the modified LEAP reactions are indicated at the top figure panels. The protein samples used in the modified LEAP reaction are indicated above the gel images. Reactions conducted with MMLV RT served as an additional control. The empty virus (no rORF2p), $H_2O$, no MMLV RT, and heat inactivated rORF2p (rORF2p_hi) served as negative controls. The black arrows indicate the expected length of the full-length modified LEAP products. At least three independent biological replicates were conducted for each condition. *(**C**) Retroviral RTs (MMLV RT and AMV RT) cannot initiate reverse transcription from the T4+3T*invT oligonucleotide primer*. MMLV RT and AMV RT could initiate reverse transcription using the T12-3'-OH oligonucleotide primer but not the T4+3T*invT primer. $H_2O$ and no RT indicate the PCR and RT reaction controls, respectively. The black arrow indicates the expected length of the full-length RT-PCR products. At least three independent biological replicates were conducted for each reaction condition. *(**D**) Exonuclease treatment of the T4+VN-3'-OH (MN440), T4+AAddC (MN488), and T4+3T*invT (MN503) oligonucleotides* (**Table S1**). Oligonucleotides were radiolabeled using [$\gamma^{32}$-P]-ATP and were incubated at 37°C in the presence (+) or absence (-) of Exonuclease I (ExoI) for the indicated time (minutes). The oligonucleotides then were separated on a denaturing polyacrylamide gel and visualized by autoradiography. At least three independent technical replicates were conducted for each reaction condition. As opposed to the T4+VN-3'-OH and T4+AAddC oligonucleotides, the T4+3T*invT oligonucleotide was resistant to exonuclease I (ExoI) digestion.

**Figure S4: Additional controls for the RT-independent ligation-based assay to monitor alt-EN activity (see Figure 3)**. *(**A**) Product detection depends on all reaction components*. The x-axis indicates the experimental condition: T4-3'-OH, the inclusion of all reaction components; T4-3'-OHnobridge, reactions lacking the bridge oligonucleotide; T4-3'-OHnoPDNA, reactions lacking the 5' phosphorylated DNA oligonucleotide; noT4-3'-OH, reactions lacking the T4-3'-OH oligonucleotide. The y-axis shows the average amount of ligated DNA. The error bars represent the SD of three technical replicates in independent experiments (see **Materials and Methods**). ***(B)*** *DNA Ligase can distinguish the 3’ ends* *of the substrate oligonucleotides.* The x-axis indicates cleavage reactions using the T4-3'-OH (MN526), T4+AAddC (MN528), and T4+3T*invT (MN527) oligonucleotides (**Table S1**). The 15 µl reactions were performed using the following buffer conditions: L1 RNPs (1.5 µl RNPbuffer, 1x cOmplete EDTA-free Protease Inhibitor Cocktail [Roche]); IP-ORF2p (1.5 µl IPbuffer, Lysis300 buffer); and rORF2p (2.0 µl rORF2pbuffer, Lysis Buffer). $H_2O$ indicates samples incubated in water. Samples lacking ligase (-Lig) were used as a negative control. The y-axis indicates the average amount of ligated DNA product (nM). The error bars represent the SD from three technical replicates in independent experiments. ***(C)*** The effect of RNA addition on rORF2p oligonucleotide cleavage. The T4+AAddC (**Table S1**: MN528, left) and T4+3T*invT (**Table S1**: MN527, right) were incubated with rORF2p or without rORF2p (Buffer). The x-axis indicates the RNA template included with rORF2p in the cleavage reaction (+23A, addition of an RNA containing 23 adenine residues [**Table S1**: MN584]; +23U, addition of an RNA containing 23 uracil residues [**Table S1**: MN585]). The y-axis indicates the average amount of ligated DNA normalized to the buffer control. Samples lacking DNA ligase (-Lig) were used as a negative control. The error bars represent the SD from three independent experiments. Statistical analysis compared with the Buffer control was performed using the Student's t-test; *P < 0.05; **P < 0.01; ***P < 0.001.

**Figure S5: RT-independent ligation-based assays to monitor alt-EN activity using L1 RNP and IP-ORF2p samples.** *(**A** and **D**) Rationale of the Assay.* The rationale of the assay is the same as that described in **Fig. 3A**. In panel A, crude RNPs were used as the source of ORF2p. In panel D, IP-ORF2p samples were used as the source of ORF2p. (***B** and **C***) *Results from the RT-independent ligation-based assay using L1 RNP samples.* Reactions were conducted using oligonucleotides the ended in either T4+AAddC (panel B) or T4+3T*invT (panel C). The X-axis indicates the L1 RNP sample mixed with the oligonucleotide in the cleavage reaction. Controls: pCEP4 (empty vector); buffer (no protein); hi, heat-inactivated; noRNase, pre-incubated without RNase; +RNase, pre-incubated with RNase A/I; -Lig, no ligase in the ligation reaction. The Y-axis indicates the average amount of ligated DNA products detected after normalization to the buffer control. The error bars represent the standard deviation from five independent experiments. A statistical analysis compared with the Buffer control was performed using the Student's t-test; *P < 0.05; **P < 0.01. (***D** and **E***) *Results from the RT-independent ligation-*

*based assay using IP-ORF2p samples.* Reactions were conducted using oligonucleotides the ended in either T4+AAddC (panel E) or T4+3T*invT (panel F). The X-axis indicates the IP-ORF2p sample used in the reaction. Controls: pCEP4 (empty vector); pAD001 (ORF2p without an epitope tag); buffer (no protein); $H_2O$ (PCR control); hi, heat-inactivated; noRNase, pre-incubated without RNase; +RNase, pre-incubated with RNase A/I; -Lig, no ligase control. The Y-axis indicates the average amount of ligated DNA products detected after normalization to the buffer control. The error bars represent the SD in four *(panel E)* or five *(panel F)* independent experiments. A statistical analysis compared with the Buffer control was performed using Student's t-test; *P < 0.05; **P < 0.01; ***P < 0.001.

**Figure S6: Modified LEAP reactions using different rORF2p protein preparations and Induro RT**. *(**A** and **B**) Flow-charts of rORF2p purification with PEI precipitation (panel A) and after the removal of IGEPAL CA-630 (panel B)*. High Five cells were infected with a baculovirus encoding a WT and mutant versions of L1.3 rORF2p containing an in-frame 3xFLAG/3xHA epitope tag at its carboxyl-terminus. After three days of incubation, the cells were lysed, and the resultant whole cell lysates were subjected to PEI precipitation (panel A). The recombinant ORF2 proteins (rORF2p) then were purified using: (**i**) anti-FLAG affinity chromatography; (**ii**) anti-HA affinity chromatography; and (**iii**) heparin column chromatography. To remove the IGEPAL CA-630 (panel B), the heparin column was washed with 20 column volumes of buffer lacking IGEPAL CA-630. The eluate then was dialyzed to obtain purified rORF2p. (see **Materials and Methods**). *(**C**) Visualization of the purified proteins*. The purified protein samples (rORF2p, rORF2p+PEI, and rORF2p_no IGEPAL) were separated on a denaturing polyacrylamide gel and visualized by silver staining. At least two independent biological replicates were conducted for each purified sample. *(**D**) Results of the modified LEAP assay.* Modified LEAP reactions were conducted using the JS+23A template RNA as described in **Fig. 4A**. The oligonucleotide primer is indicated at the top of each gel. The protein samples prepared in the absence (no PEI) or presence (+PEI) of a polyethyleneimine precipitation step are indicated above each gel image. The protein used in each modified LEAP reaction is indicated above the gel image. Buffer (no protein control), empty virus (no protein control), $H_2O$ (PCR control). Marker, molecular weight standards in base pairs (bp). The black arrow indicates the size of the expected full-length product. At least three independent biological replicates were conducted for each reaction condition. *(**E** and **F**) A commercially available thermostable group II intron-encoded protein has alt-EN activity.* Modified LEAP reactions were conducted using the JS+23A template RNA performed as described in **Fig. 4D**. The protein used in each modified LEAP reaction is indicated above the gel image. The oligonucleotide primer is indicated at the top of each gel. In panel E, modified LEAP reactions were performed at 37°C. In panel F, the reactions containing the Induro RT were performed at 55°C. Controls: no primer (RT reaction negative control); no protein (no protein included in reaction); $H_2O$ (PCR control); heat inactivated Induro RT (negative control); proteinase K (ProK) inactivated Induro RT (negative control); rORF2p (positive control); MMLV RT (retroviral reverse transcriptase control). Marker, molecular weight standards in base pairs (bp). At least three independent replicates were conducted for each experimental condition. *(**G**) Characterization of Products from the modified LEAP reaction*. The products from **Fig. 4D** and **Fig. S6E** were characterized by DNA sequencing. The labeling is like that described in **Fig. 1C**, **Fig. 1G**, **and Fig. S2G**. *(**H**) Characterization of modified LEAP products using the T4+VN-3'-OH primer from **Fig. 4D** and **Fig. S6E***. Labeling is like that in **Fig. S2H**. The "V" and "N" nucleotides in the modified LEAP reaction products are indicated in the Table. Note: Induro RT can extend primer/template complexes that lack perfect complementarity (41,54).

#### Supplemental Dataset

The Supplemental Dataset detailing the results from our product characterization assays are indicated in separate tabs and can be downloaded using the following link:

https://www.dropbox.com/scl/fi/9dz5vm8ool952nrnfdy45/FINAL-SUPPLEMENTAL-DATASET-Nakamura-et-al.-1-17-2026.xlsx?rlkey=wdj8u75bmppbkun0rw4e1gjfa&st=c82hcn2i&dl=0

**Table S1: Oligonucleotides used in this study.**

| Name | Sequence |
|---|---|
| MN056 | 5'-TAATACGACTCACTATAGGGAATTGAACAATGAGATCACATG-3' |
| MN057 | 5'-TTTTTTTTTTTTTTTTTTTTTTTTATTATACTCTAAGTTTTAGGGTACATG-3' |
| MN058 | 5'-ATTATACTCTAAGTTTTAGGGTACATG-3' |
| MN085 | 5'-GCGAGCACAGAATTAATACGACT-3' |
| MN439 | 5'-GCGAGCACAGAATTAATACGACTCACTATAGGATCTTTT-3' |
| MN440 | 5'-GCGAGCACAGAATTAATACGACTCACTATAGGATCTTTTVN-3' |
| MN462 | 5'-GACTTTCCACACCCTAACTG-3' |
| MN468 | 5'-GTTGTTAACTTGTTTATTGCAGCTTATAATGGTTAC-3' |
| MN469 | 5'-TTTTTTTTTTTTTTTTTTTTTTTTGTTGTTAACTTGTTTATTGCAGCTTATAATGGTTAC-3' |
| MN470 | 5'-TAATACGACTCACTATAGGGGAGCCTGGGGACTT-3' |
| MN485 | 5'-GGGTTCGAAATCGATAAGCTTGGATCCAGAC-3' |
| MN488 | 5'-GCGAGCACAGAATTAATACGACTCACTATAGGTTTTAAddC-3' |
| MN503 | 5'-GCGAGCACAGAATTAATACGACTCACTATAGGATCTTTTT*T*T*invT-3' |
| MN526 | 5'biotin-18atomhexa-ethyleneglycolspacer-GCGAGCACAGAATTAATACGACTCACTATAGGATCTTTT-3' |
| MN527 | 5'biotin-18atomhexa-ethyleneglycolspacer-GCGAGCACAGAATTAATACGACTCACTATAGGATCTTTTT*T*T*invT-3' |
| MN528 | 5'biotin-18atomhexa-ethyleneglycolspacer-GCGAGCACAGAATTAATACGACTCACTATAGGATCTTTTAAddC-3' |
| MN544 | 5'Phosphate-GCTTCCGTCTCTGTCCTTGTGTGCAGGAGANNNNNCGATGGAGGCGTTGGAGCTGGAGCTGGAAGAAGTGGAGTCCCAGATCCGCGCGCTGGTGGTAAGACGGTCGCGGCTACGGGAACGGCTTCTAGCCGTACCTAATGCTAAGGCCGTCTCATCACCTAAGGTACGTGGAAATTACAACCACATCATTCCCTCTACCT-3' |
| MN546 | 5'-GACAGAGACGGAAGCAAAAGATCCTATAGTGAGTCinvdT-3' |
| MN547 | 5'-GCGAGCACAGAATTAAT-3' |
| MN549 | 5'-GAATGATGTGGTTGTAATTT-3' |
| MN551 | 5'-FAM-TCCAGCTCC-ZEN-AACGCCTCCATC-IowaBlakFQ-3' |
| MN552 | 5'-ACGACTCACTATAGGATCTTTTGC-3' |
| MN553 | 5'-CTAGAAGCCGTTCCCGTAG-3' |
| MN584 | 5'-rArArArArArArArArArArArArArArArArArArArArA-3' |
| MN585 | 5'-rUrUrUrUrUrUrUrUrUrUrUrUrUrUrUrUrUrUrUrU-3' |
| MN746 | 5'-GCGAGCACAGAATTAATACGACTCACTATAGGATCTTTTT*T*T*ddC-3' |
| ORF2M | 5'-GAGATCACATGGACACAGGAAGGG-3' |
| T12 | 5′-GCGAGCACAGAATTAATACGACTCACTATAGGTTTTTTTTTTTT-3′ |
| 4228 | 5′-GCGAGCACAGAATTAATACGACTCACTATAGGATCTTTTTTT-3′ |

### Figure S1

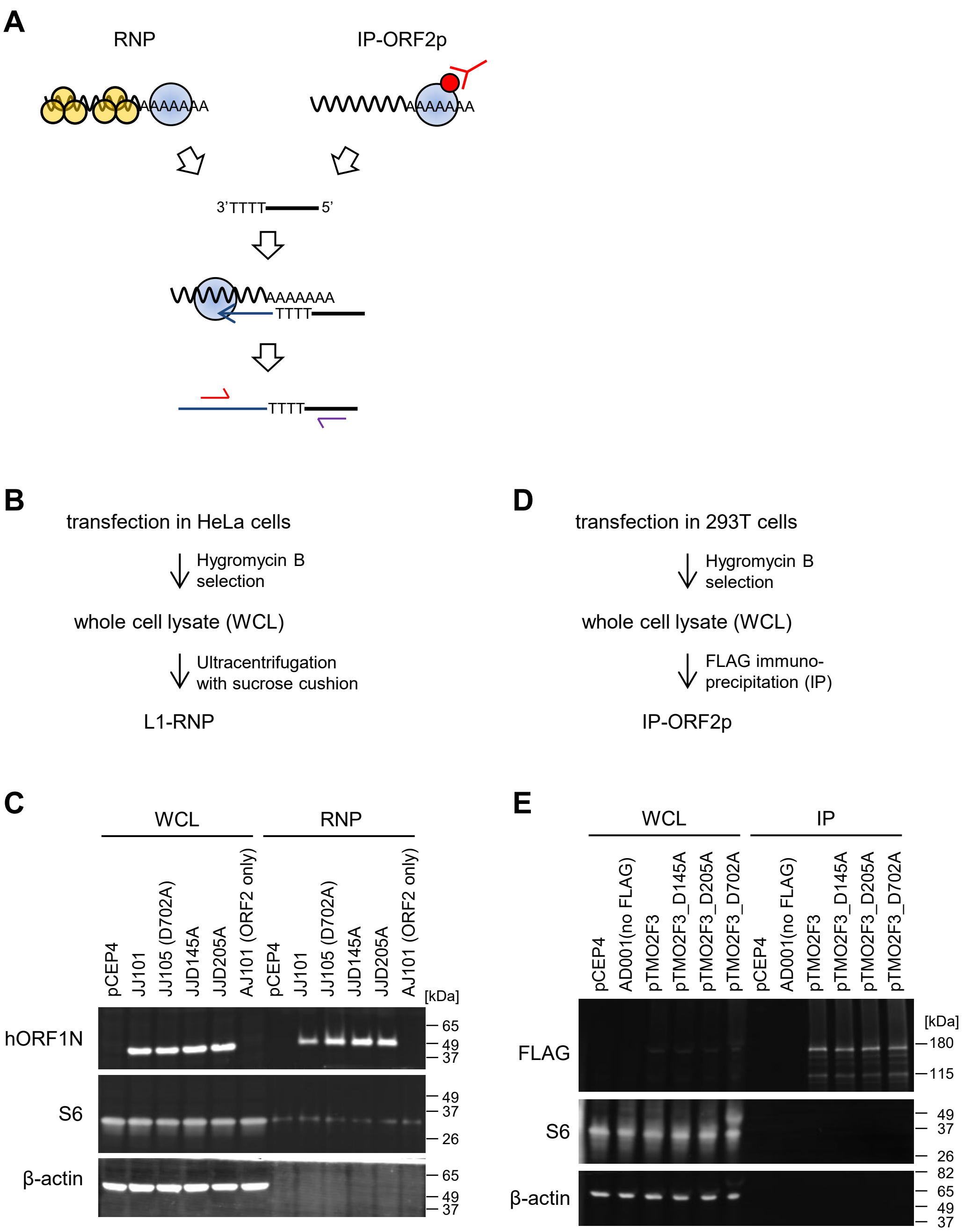
A
RNP
IP-ORF2p
AAAAAAA
3'TTTT
5'
TTTT
B
transfection in HeLa cells
Hygromycin B selection
whole cell lysate (WCL)
Ultracentrifugation with sucrose cushion
L1-RNP
D
transfection in 293T cells
Hygromycin B selection
whole cell lysate (WCL)
FLAG immuno-precipitation (IP)
IP-ORF2p
C
WCL
RNP
pCEP4
JJ101
JJ105 (D702A)
JJD145A
JJD205A
AJ101 (ORF2 only)
[kDa]
hORF1N
S6
β-actin
65
49
37
26
E
WCL
IP
pCEP4
AD001(no FLAG)
pTMO2F3
pTMO2F3_D145A
pTMO2F3_D205A
pTMO2F3_D702A
FLAG
180
115
82

### Figure S2

A

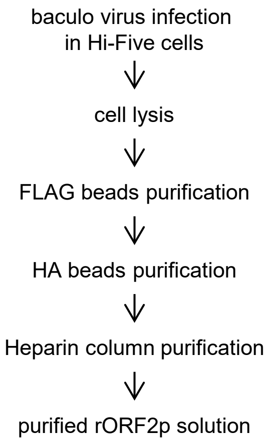


B

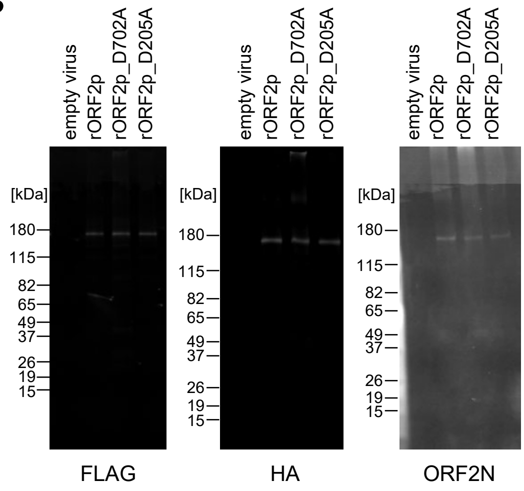


C

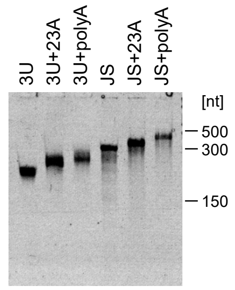


D

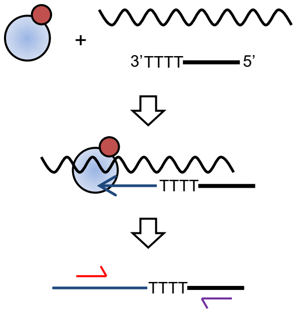


E

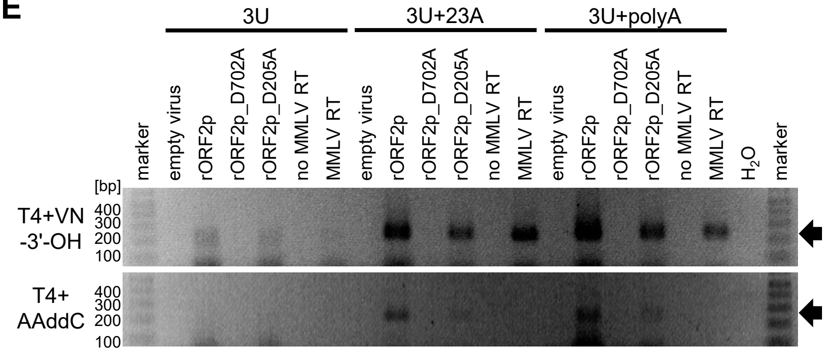


F

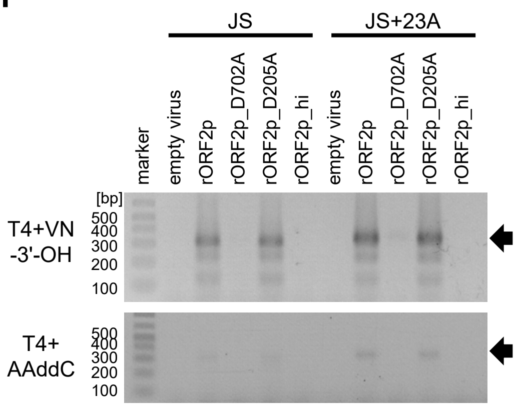


G

| | | T4+VN-3'-OH | | T4AAddC | | |
|---|---|---|---|---|---|---|
| | | T1-T4 | T4VN | T1-T4 | T4A(A) | T4AAC |
| rORF2p | 3U | 0/10 | 10/10 | n.d. | n.d. | n.d. |
| | 3U+23A | 0/10 | 10/10 | 9/16 | 7/16 | 0/16 |
| | 3U+polyA | 1/11 | 10/11 | 5/10 | 5/10 | 0/10 |
| | JS | 0/10 | 10/10 | n.d. | n.d. | n.d. |
| | JS+23A | 0/10 | 10/10 | 4/10 | 6/10 | 0/10 |
| | JS+polyA | 0/10 | 10/10 | 8/10 | 2/10 | 0/10 |
| rORF2p_D205A | 3U | 0/10 | 10/10 | n.d. | n.d. | n.d. |
| | 3U+23A | 0/10 | 10/10 | 8/14 | 6/14 | 0/14 |
| | 3U+polyA | 0/10 | 10/10 | 3/10 | 7/10 | 0/10 |
| | JS | 0/11 | 11/11 | n.d. | n.d. | n.d. |
| | JS+23A | 0/10 | 10/10 | 6/10 | 4/10 | 0/10 |
| | JS+polyA | 0/10 | 10/10 | 7/11 | 4/11 | 0/10 |
| MMLV-RT | 3U | 0/12 | 12/12 | n.d. | n.d. | n.d. |
| | 3U+23A | 0/10 | 10/10 | n.d. | n.d. | n.d. |
| | 3U+polyA | 0/10 | 10/10 | n.d. | n.d. | n.d. |
| | JS | 0/11 | 10/11 | n.d. | n.d. | n.d. |
| | JS+23A | 0/10 | 10/10 | n.d. | n.d. | n.d. |
| | JS+polyA | 0/11 | 11/11 | n.d. | n.d. | n.d. |

H

| rORF2p | | N: a | t | g | c |
|---|---|---|---|---|---|
| V | a | 0 | 2 | 0 | 2 |
| | c | 0 | 3 | 0 | 0 |
| | g | 1 | 4 | 2 | 1 |

| rORF2p _D205A | | N: a | t | g | c |
|---|---|---|---|---|---|
| V | a | 0 | 6 | 2 | 1 |
| | c | 0 | 0 | 1 | 1 |
| | g | 1 | 0 | 0 | 0 |

| MMLV -RT | | N: a | t | g | c |
|---|---|---|---|---|---|
| V | a | 0 | 0 | 0 | 0 |
| | c | 0 | 0 | 0 | 0 |
| | g | 0 | 8 | 0 | 0 |

### Figure S3

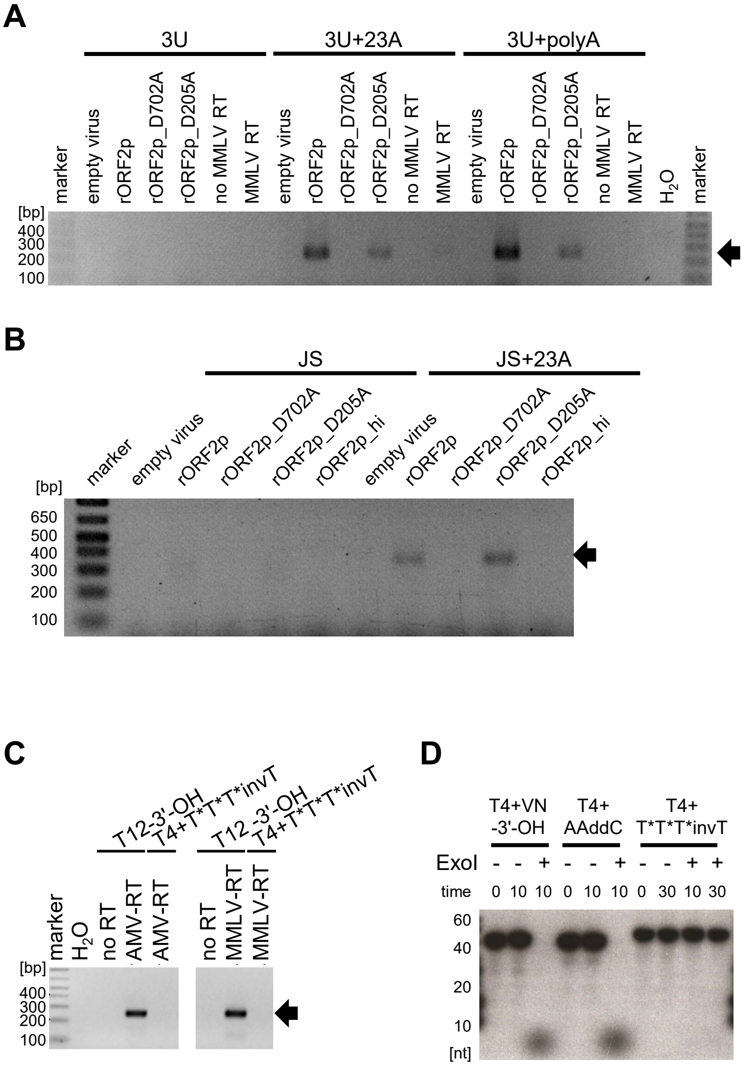

A
3U
3U+23A
3U+polyA
marker
empty virus
rORF2p
rORF2p_D702A
rORF2p_D205A
no MMLV RT
MMLV RT
$H_2O$
[bp]
400
300
200
100
B
JS
JS+23A
rORF2p_hi
650
500
C
T12-3'-OH
T4+T*T*T*invT
no RT
AMV-RT
MMLV-RT
D
T4+VN
-3'-OH
T4+
AAddC
T*T*T*invT
ExoI
time
60
40
20
10
[nt]

### Figure S4

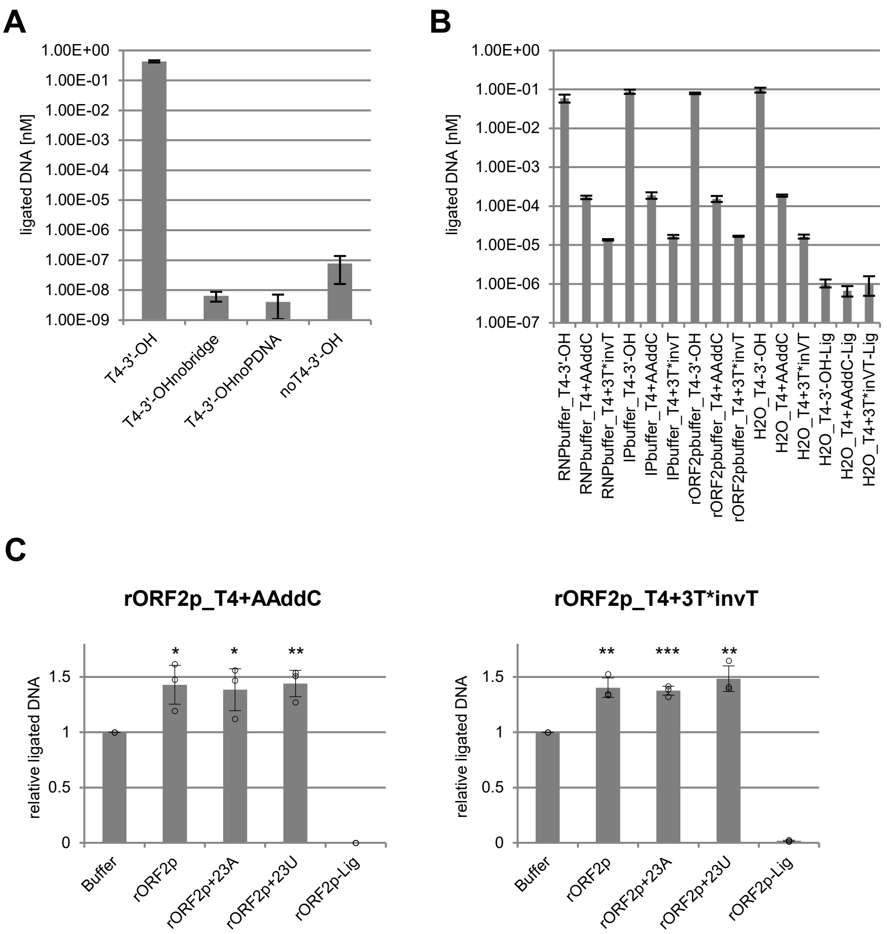
A
1.00E+00
1.00E-01
1.00E-02
1.00E-03
1.00E-04
1.00E-05
1.00E-06
1.00E-07
1.00E-08
1.00E-09
ligated DNA [nM]
T4-3'-OH
T4-3'-OHnobridge
T4-3'-OHnoPDNA
noT4-3'-OH
B
1.00E+00
1.00E-01
1.00E-02
1.00E-03
1.00E-04
1.00E-05
1.00E-06
1.00E-07
ligated DNA [nM]
RNPbuffer_T4-3'-OH
RNPbuffer_T4+AAddC
RNPbuffer_T4+3T*invT
IPbuffer_T4-3'-OH
IPbuffer_T4+AAddC
IPbuffer_T4+3T*invT
rORF2pbuffer_T4-3'-OH
rORF2pbuffer_T4+AAddC
rORF2pbuffer_T4+3T*invT
H2O_T4-3'-OH
H2O_T4+AAddC
H2O_T4+3T*inVT
H2O_T4-3'-OH-Lig
H2O_T4+AAddC-Lig
H2O_T4+3T*inVT-Lig
C
rORF2p_T4+AAddC
relative ligated DNA
0
0.5
1
1.5
*
*
**
Buffer
rORF2p
rORF2p+23A
rORF2p+23U
rORF2p-Lig
rORF2p_T4+3T*invT
relative ligated DNA
0
0.5
1
1.5
**
***
**
Buffer
rORF2p
rORF2p+23A
rORF2p+23U
rORF2p-Lig

### Figure S5

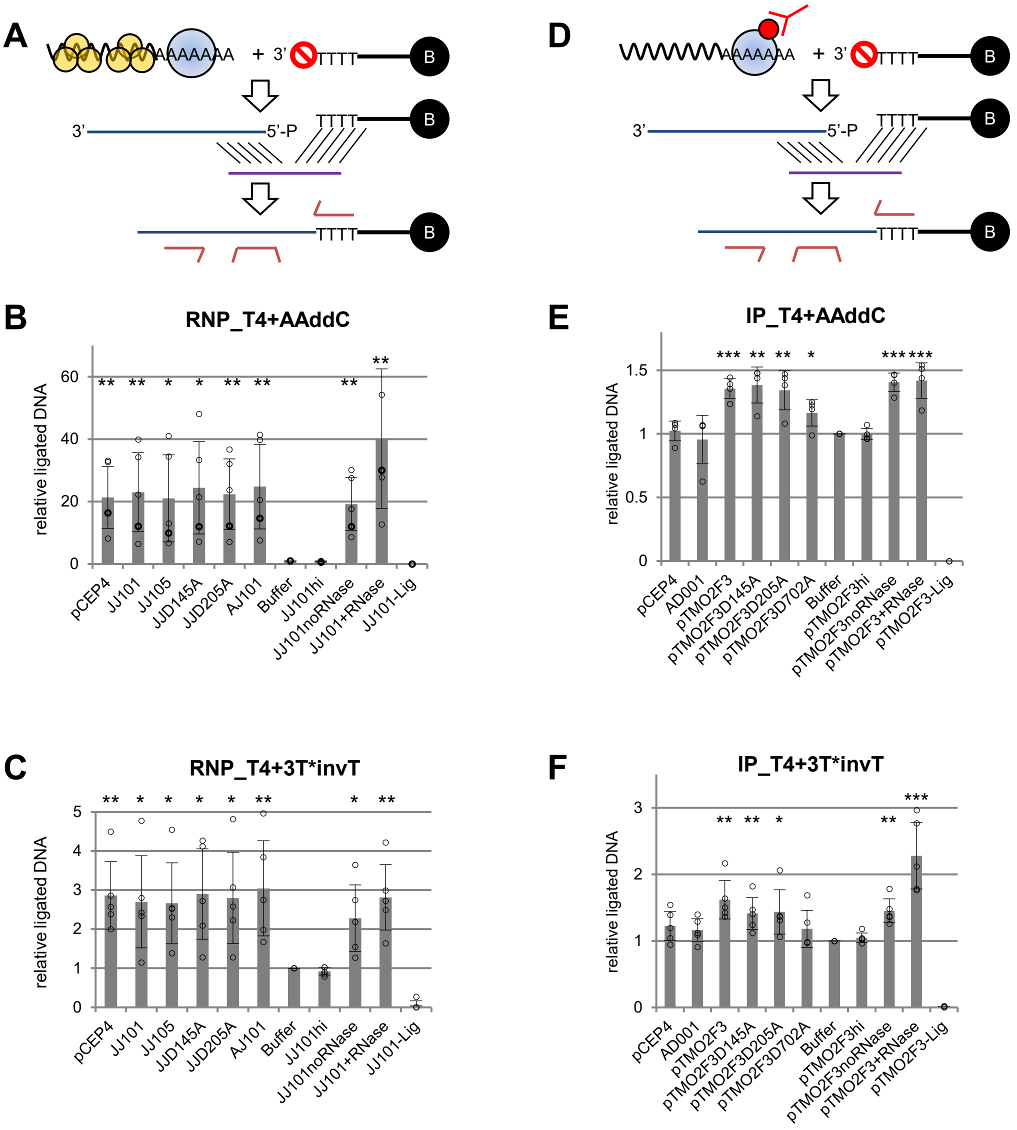

A
AAAAAAA
+ 3'
TTTT
B
3'
5'-P
D
B
RNP_T4+AAddC
relative ligated DNA
pCEP4
JJ101
JJ105
JJD145A
JJD205A
AJ101
Buffer
JJ101hi
JJ101noRNase
JJ101+RNase
JJ101-Lig
C
RNP_T4+3T*invT
E
IP_T4+AAddC
AD001
pTMO2F3
pTMO2F3D145A
pTMO2F3D205A
pTMO2F3D702A
pTMO2F3hi
pTMO2F3noRNase
pTMO2F3+RNase
pTMO2F3-Lig
F
IP_T4+3T*invT

### Figure S6

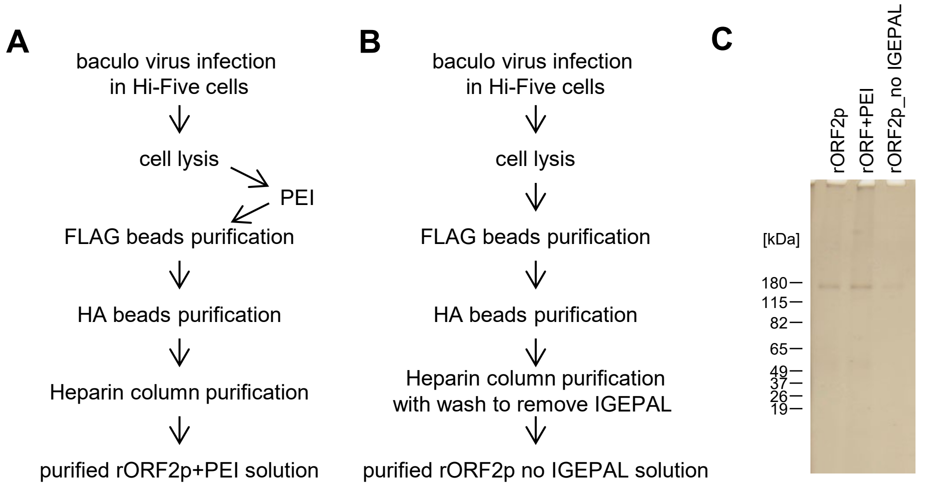


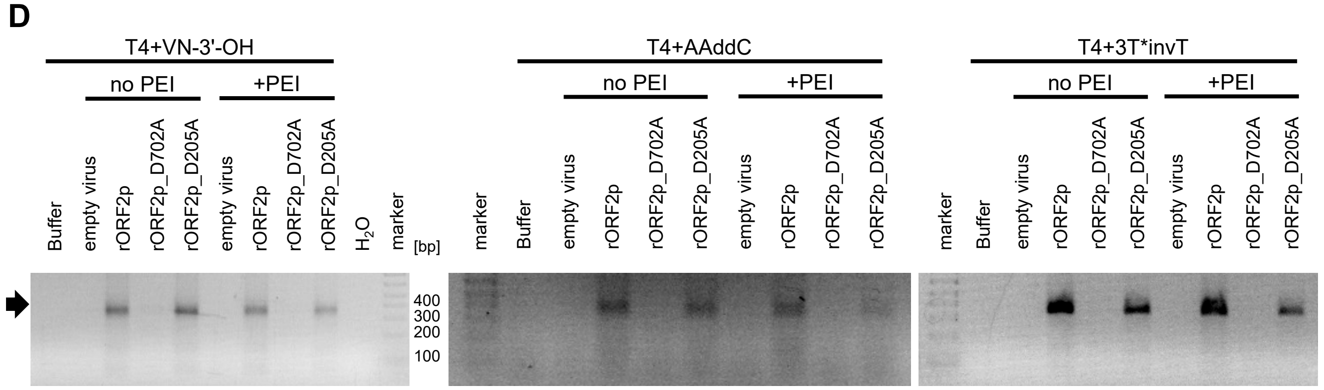


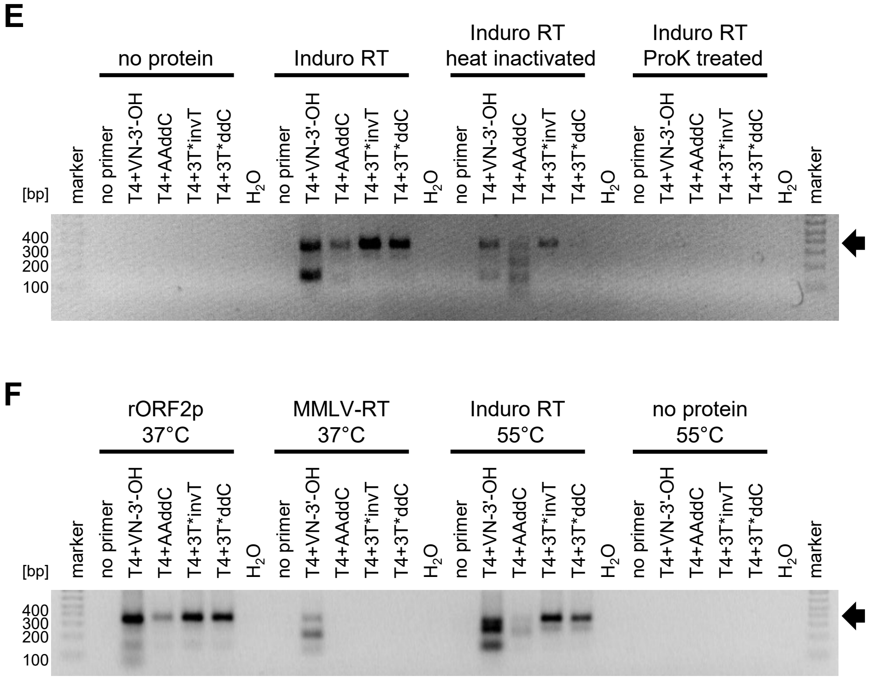


**G**

| | T4+VN-3'-OH | | T4+AAddC | | |
|---|---|---|---|---|---|
| | T1-4 | T4VN | T1-T4 | T4A(A) | T4AAC |
| Induro RT | 1/30 | 30/31 | 8/30 | 21/30 | 1/30 |

**H**

| Induro RT | | N | | | |
|---|---|---|---|---|---|
| | | a | t | g | c |
| V | a | 0 | 0 | 0 | 2 |
| V | c | 1 | 1 | 0 | 0 |
| V | g | 1 | 4 | 4 | 1 |
